# FRUCTOSE ACTIVATES A STRESS RESPONSE SHARED BY METHYLGLYOXAL AND HYDROGEN PEROXIDE IN *STREPTOCOCCUS MUTANS*

**DOI:** 10.1101/2024.10.26.620100

**Authors:** Alejandro R. Walker, Danniel N. Pham, Payam Noeparvar, Alexandra M. Peterson, Marissa K. Lipp, José A. Lemos, Lin Zeng

## Abstract

Fructose catabolism by *Streptococcus mutans* is initiated by three PTS transporters yielding fructose-1-phoshate (F-1-P) or fructose-6-phosphate. Deletion of one such F-1-P-generating PTS, *fruI*, was shown to reduce the cariogenicity of *S. mutans* in rats fed a high-sucrose diet. Moreover, a recent study linked fructose metabolism in *S. mutans* to a reactive electrophile species (RES) methylglyoxal. Here, we conducted a comparative transcriptomic analysis of *S. mutans* treated briefly with 50 mM fructose, 50 mM glucose, 5 mM methylglyoxal, or 0.5 mM hydrogen peroxide (H_2_O_2_). The results revealed a striking overlap between the fructose and methylglyoxal transcriptomes, totaling 176 genes, 61 of which were also shared with the H_2_O_2_ transcriptome. This core of 61 genes encompassed many of the same pathways affected by exposure to low pH or zinc intoxication. Consistent with these findings, fructose negatively impacted metal homeostasis of a mutant deficient in zinc expulsion and the growth of a mutant of the major oxidative stress regulator SpxA1. Importantly, fructose metabolism lowered culture pH at a faster pace, allowed better survival under acidic and nutrient-depleted conditions, and enhanced the competitiveness of *S. mutans* against *Streptococcus sanguinis*, although a moderated level of F-1-P might further boost some of these benefits. Conversely, several commensal streptococcal species displayed a greater sensitivity to fructose that may negatively affect their persistence and competitiveness in dental biofilm. In conclusion, fructose metabolism is integrated into the stress core of *S. mutans* and regulates critical functions required for survival and its ability to induce dysbiosis in the oral cavity.

**Importance.:** Fructose is a common monosaccharide in the biosphere, yet its overconsumption has been linked to various health problems in humans including insulin resistance, obesity, diabetes, non-alcoholic liver diseases, and even cancer. These effects are in large part attributable to the unique biochemical characteristics and metabolic responses associated with the degradation of fructose. Yet, an understanding of the effects of fructose on the physiology of bacteria and its implications to the human microbiome is severely lacking. Here we performed a series of analyses on the gene regulation of a dental pathogen *Streptococcus mutans* by exposing it to fructose and other important stress agents. Further supported by growth, persistence, and competition assays, our findings revealed the ability of fructose to activate a set of stress-related functions that may prove critical to the ability of the bacterium to persist and cause diseases both within and without of the oral cavity.

## Introduction

Disruption in microbial homeostasis in human microbiomes, a condition termed dysbiosis, has been associated with the development of many diseases that range from GI tract disorders such as inflammatory bowel disease and pseudomembranous colitis to oral infectious diseases such as dental caries, candidiasis, and periodontitis (1, 2). Carbohydrate metabolism plays a vital role in oral health by contributing to the biochemistry and ecology of oral microbiota. It is widely understood that overconsumption of refined carbohydrates is a defining factor in the development of dysbiosis through acid production and establishment of an acidogenic and aciduric dental biofilm (3–6). Emerging evidence also supports the co-occurrence of dental caries and periodontal diseases, pointing to potentially shared mechanisms in underlying etiology (7–9). With the food industry shifting heavily toward the use of fructose and fructose-releasing sugars in the past several decades and the concurrent pandemic of metabolic syndrome, the impact of dietary fructose to oral and systemic health demands urgent attention (10).

Fructose is a ubiquitous nutrient widely found in fruits, yet overconsumption of fructose is often associated with adverse health outcomes in humans that include insulin resistance, obesity, liver diseases, systemic inflammation, type-2 diabetes, cogitative decline, and even cancer (11–15). According to the “fructose survival hypothesis”, fructose triggers a conserved cellular response in mammalian cells that manifests as mitochondrial oxidative stress and reduced cellular ATP levels, accompanied by behavioral changes that includes hunger, thirst, foraging, and weight gain (16). While this response could have been limited and beneficial to animals and our human ancestors, the negative effects of fructose have been greatly exacerbated by its widespread use in Western diet.

Streptococci are the most abundant commensal species colonizing oral surfaces, and most of them rely on carbohydrate fermentation, via the Embden–Meyerhof–Parnas (EMP) pathway of glycolysis, to produce energy and precursors for various biogenesis processes (3). Multiple fructose-transporting mechanisms exist in streptococci, each harboring a fructose: phosphotransferase (PTS) permease that generates either fructose-1-phosphate (F-1-P) or fructose-6-phosphate (F-6-P), which is further phosphorylated into F-1,6-bP by their respective phosphofructokinases (17). Studies so far have indicated that F-1-P metabolism contributes to *in vitro* fitness, biofilm development, and virulence for pathogenic species such as *Streptococcus pyogenes, Streptococcus agalactiae,* and *Streptococcus gordonii* (18–22). For the cariogenic pathobiont *Streptococcus mutans*, the primary fructose pathway is encoded by the *fruRKI* operon, encoding a negative regulator FruR, a PTS permease (FruI) that yields F-1-P, and a F-1-P kinase (FruK) (17). Deletion of *fruI* in *S. mutans* UA159 resulted in lower levels, though not complete lack of F-1-P when exposed to fructose (23) and reduced the ability of the bacterium to colonize and induce dentin caries in rats fed a high-sucrose diet (24). Based on genetic studies, expression of *fruRKI,* and its homologue *fruRBA* in related streptococci, is inducible by F-1-P which acts allosterically on FruR (17, 18, 25). Another fructose-PTS (EII^Lev^), encoded by *levDEFG,* exists in some of these oral streptococci, including *S. mutans, Streptococcus sanguinis* and *S. gordonii*, which likely generates F-6-P and requires a four-component system LevQRST for induction in response to extracellular fructose (26). *S. mutans* harbors yet another fructose-PTS (*fruCD*) that is also predicted to generate F-1-P, although its contribution to the overall fructose transport appears to be minor (17). Overall, the F-1-P pathway encoded by *fruRBA/fruRKI* operon is conserved in most Gram-positive bacteria encompassing *Bacillus, Enterococcus, Lactococcus, Lactobacillus, Clostridium, Listeria,* and most *Streptococcus* species (27–29). Moreover, *S. mutans, S. gordonii,* and *S. sanguinis* also harbor another fructose-related operon, *sppRA.* SppA has been characterized as a hexose-phosphate phosphohydrolase that specializes in dephosphorylating F-1-P, and F-6-P to a lesser extent; and SppR is a negative auto-regulator whose activity is allosterically regulated by F-1-P (23).

A recent study involving fructose centered on the molecular mechanisms required for oral streptococci to degrade a group of highly reactive electrophile species (RES), including methylglyoxal and glyoxal (30), which are created by all life as byproducts of glycolysis and serve as important immune effectors (31, 32). It has been reported that *S. mutans* is significantly more resistant to methylglyoxal than several commensal streptococci including *S. gordonii, S. sanguinis,* and members of the mitis group (30). A zinc-containing protein GloA2 (SMU.1112c), predicted to be a paralogue to the main methylglyoxal-degrading enzyme glyoxalase A (LguL or GloA) based on its sequence and crystal structure, was identified in all oral streptococci examined so far (30). The *gloA2*-null mutants of both *S. mutans* and *S. sanguinis* displayed increased chaining and autolysis upon treatment by fructose which resembled the effect of exposure to methylglyoxal, including its reversal by treatment with glutathione (GSH) which is known to scavenge methylglyoxal before degradation by LguL (32). Interestingly, the expression of *lguL* was inducible by growth on fructose, and the fructose-sensitivity of *gloA2* mutants was largely dependent on the integrity of the F-1-P-generating PTS, FruI (30). Herein we further examined fructose-dependent gene regulation and physiology in relation to streptococcal tolerance to important stressors including methylglyoxal and hydrogen peroxide. Our findings revealed the integration of fructose-mediated response with general stress response experienced during bacteria-host and inter-microbial interactions.

## Results

### Comparative RNA-seq analyses identified a stress core shared by methylglyoxal, fructose, and H_2_O_2_

We previously performed an RNA-seq analysis on *S. mutans* cultured to the exponential phase with 0.5% fructose (27.7 mM) in the medium (33). The strain used in this study was a mutant deficient in several enzymes capable of degrading sucrose and other carbohydrate polymers (34), and was later revealed to contain a spontaneous mutation in a major peroxide-sensing regulator PerR (35). While this work provided us important information regarding the state of the bacterium after growing on fructose, these genetic deviations are of concern and the study could not capture the response the cells present immediately after fructose intake, as metabolic regulation tends to return to equilibrium after initial perturbance. To mimic the physiological impact to bacteria when humans consume a large dose of fructose in food or drinks, we cultured *S. mutans* strain UA159 (from ATCC) in a glucose-based medium to the exponential phase and then treated the cells with 50 mM fructose for 30 min, which were immediately processed for RNA-seq analysis in comparison to untreated controls. Considering the similarity between glucose and fructose, the same UA159 cultures were treated with 50 mM glucose in the same manner and analyzed as an additional control. These concentrations of carbohydrates represent the physiological levels the microbiota likely encounter periodically in oral cavity. The experiment was also repeated using the same bacteria after treatment with 5 mM methylglyoxal for 30 min (30). In addition, we obtained and reanalyzed the datasets of a similar RNA-seq study performed on UA159 cultures that were prepared in a glucose-based medium and treated with 0.5 mM H_2_O_2_ for 5 min (36) (see Fig. 1 for volcano plots, Fig. 2 for Venn diagrams, Fig. S1 for principal component analyses, Fig. S2 for Gene Ontology term analyses, and supplemental Tables S1 to S3 for other details).

**Fig. 1.**
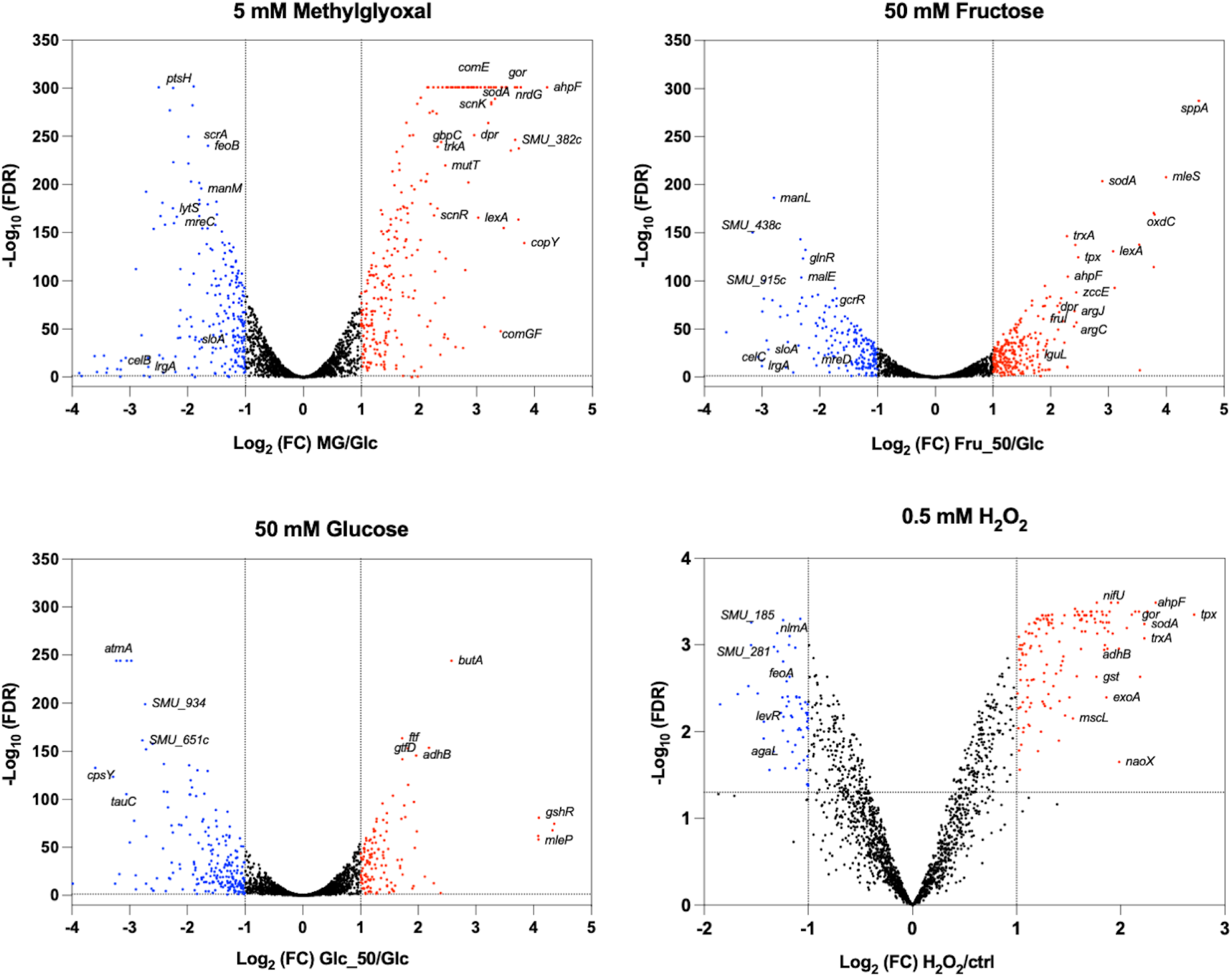
Volcano plots illustrating transcriptomes of four treatments. *S. mutans* UA159 was grown to the exponential phase in a synthetic FMC medium containing 20 mM glucose. The samples were treated for 30 min with 5 mM methylglyoxal, 50 mM fructose, or 50 mM glucose before harvesting for mRNA sequencing. Differential expression analyses were performed against untreated cells using edgeR, with cutoffs for foldchange >2 and FDR <0.05. Upregulated genes are shown in red, and downregulated ones in blue. Results of treatment by H_2_O_2_ (0.5 mM, 5 min) were from an independent study (36).

**Fig. 2.**
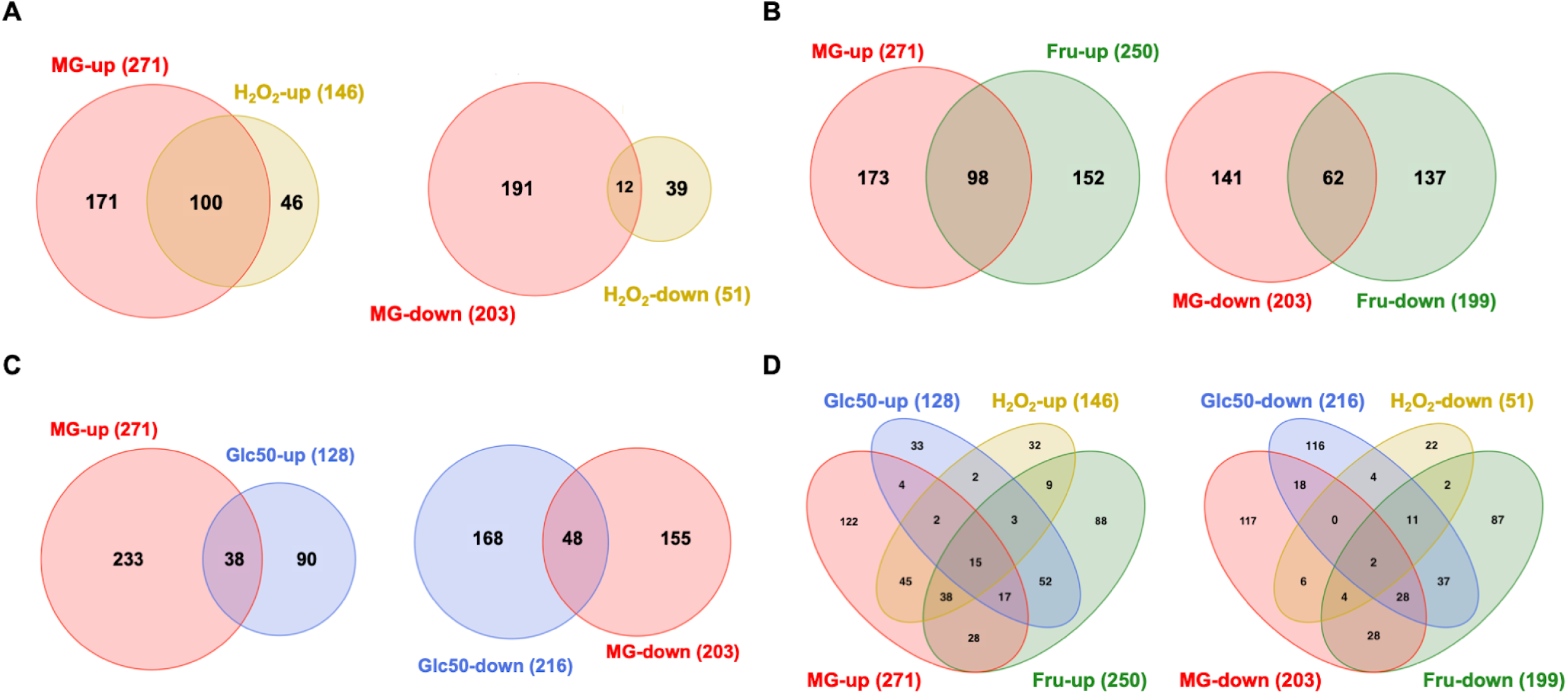
Transcriptomic overlaps among treatments of methylglyoxal, fructose, glucose and H_2_O_2._ The Venn diagrams show the numbers of shared and unique genes in the genome of UA159, separated into groups of upregulated and downregulated, among treatments of (A) 5 mM methylglyoxal (MG) and 0.5 mM H_2_O_2_, (B) MG and 50 mM fructose (Fru), (C) MG and 50 mM glucose (Glc50), and (D) all four treatments.

#### Methylglyoxal

Treatment of UA159 with 5 mM of methylglyoxal differentially affected the expression of a total of 474 genes when compared to the mock-treated control group, with 271 being up-regulated and 203 being down-regulated (Fig. 2, Fig. S2). Methylglyoxal is known to react with the reduced form of glutathione (GSH), which is interconnected with other reductive potentials such as NADPH and ascorbate (37). Treatment with methylglyoxal resulted primarily in several effects similar to oxidative stress response (36): increased expression of genes involved in thiol homeostasis including thioredoxins *trxAB*, GSH synthetase *gshAB* (38), glutathione reductase *gor,* and glutathione S-transferase *gst*; increased expression of the iron-sulfur (Fe-S) cluster biosynthesis system *sufCDSUB* (39), the ascorbate metabolic cluster (SMU.271 to SMU.275) (40), the reactive oxygen species (ROS) scavenging genes *sodA*, *ahpCF*, and *tpx*, and genes whose products are involved in metal homeostasis: copper exporter operon *copYAZ*, and iron-sequestering gene *dpr*. Also upregulated were several transcription regulators known to be involved in stress management: *relS*, *perR*, *phoR/ycbL*, *scnKR*, *rcrRPQ*, SMU.1097c, *vicKX*, *comR*, *comDE*, and *lexA*. Likely as the need to restore the GSH/GSSG balance, treatment of methylglyoxal upregulated several putative aldo/keto reductases: SMU.382c and SMU.383c, SMU.677 to SMU.681, SMU.1040c, and SMU.1602 residing immediately upstream of *lguL* which encodes the major methylglyoxal-degrading enzyme LguL/GloA (both up by 3∼4-fold). It is worth noting that most of these aldo/keto reductases, other than *lguL*, are absent in related commensal streptococci. Also related to improved reducing capacity was upregulation of several putative ribonucleotide reductases (41, 42): SMU.667 to SMU.669c, SMU.991, and SMU.2070 to SMU.2074 (*nrdG*).

Interestingly, methylglyoxal treatment reduced the expression of iron/manganese transporter *sloAB* (not *sloC*) and cognate *sloR* regulator. It also reduced the expression of ferrous ion transporter genes *feoAB*. These changes were consistent with the purported ability of GSH to buffer intracellular metal ions (43), as a release of these ions would likely increase the expression of metal exporters (*copYA*) but reduce that of metal importers (*sloABC* and *feoAB*). Another likely effect of methylglyoxal exposure is its reaction with intracellular amino acids such as arginine and lysine. For this, we observed increased expression of amino acid biosynthesis/transport operons, *argCJBD* for arginine and *livFGMHK* for branched-chain amino acids. As a general response to stressful stimuli, we observed elevated expression of genes required for DNA repair (*exoA*, *end3*, *mutY*, and *uvrA*) and protein or peptide degradation (*clpC, clpE, pepO*, and *pepP*). Conversely, there were also several stress management genes that showed reduced expression: *mreCD*, *irvR*, *hrcA, grpE,* and *dnaK*. Most ribosomal proteins and the translation initiation factor InfC also had lower mRNA levels, indicative of reduced protein synthesis activity. There was likely a shift toward a more efficient, mixed-acid fermentation as we saw greater expression by acetoin metabolic operon (*adhABCD-lplA*), the related alpha-acetolactate metabolic genes *alsS* and *aldB*, NADH oxidase (*nox*), and a pyruvate-formate lyase activating enzyme *pflA*. At the same time, the genes encoding the two major enzymes of the PTS, *ptsH* and *ptsI*, along with the transport and metabolic pathways for mannitol (EII^Mtl^, SMU.1184c and SMU.1185), cellobiose (EII^Cel^, SMU.1596 to SMU.1600), sucrose (EII^Scr^, SMU.1841 to SMU.1844), and glucose (EII^Man^, SMU.1877 to SMU.1879) all showed significantly reduced expression.

#### Comparative analysis of methylglyoxal regulon with H_2_O_2_-mediated stress response

During the analysis of the methylglyoxal-induced transcriptomic shift, it was immediately apparent that many of these genes were also identified by a previous study on peroxide stress response of *S. mutans* UA159 (36). A closer inspection of the peroxide stress transcriptome identified a total of 115 genes (Supplemental Tables S1 and S4, Fig. 2A) as being shared by methylglyoxal and H_2_O_2_, indicative of the common bacterial mechanisms needed to deal with these two stressors. These aforementioned functions included ROS scavenging, thiol and metal homeostasis, DNA repair and protein processing, and energy metabolism. Out of these 115 genes, 100 showed upregulation and 12 were downregulated; only 3 showed different polarity of change. Important distinctions were also noted which encompassed 359 methylglyoxal-specific genes that were missing in H_2_O_2_ regulon, and conversely 79 H_2_O_2_-responsive genes absent in the methylglyoxal regulon (Supplemental Table S4). Briefly, absent in H_2_O_2_-induced response included the reduced expression of the PTS genes, namely *ptsH, ptsI*, and EII^Mtl^, EII^Cel^, EII^Scr^, and EII^Man^ operons. Also absent were genes for ribosomal proteins, *lrgAB*-*lytST*, *argCJBD* and agmatine deiminase operon (44), several putative aldo/keto reductase clusters, potassium transporters *trkAB*, the *nrd* operon SMU.2070 to SMU.2074, *mreCD*, *hrcA-grpE-dnaK*, and transcription regulators including *mleR, relS*, *phoR/ycbL*, *nagR*, *irvR, vicKX, comX,* and *ahrC*. Missing from the methylglyoxal regulon were some of the H_2_O_2_-induced genes, including mutacin genes (*nlmABCD* and *nlmTE*), *sloC*, *liaSR, levQRST, relA,* and the zinc import operon *adcBCR.* Interestingly, while methylglyoxal induced expression of most of the competence regulators, *comCDE, comR*, and *rcrR* (45), and the late-competence operon *comYA*-*YI* (46), H_2_O_2_ reduced expression of two important *nlm* operons, one of which (*nlmABCD*) requires an active *comDE* circuit for expression (47).

#### Significant overlap between fructose, methylglyoxal, and H_2_O_2_-mediated responses

Treatment of UA159 with 50 mM fructose for 30 min differentially affected the expression of a total of 449 genes, with 250 being up-regulated and 199 being down-regulated (Fig. 2B, Fig. S2, and Table S3). Though not a critical point of this study, these findings represent a striking difference in gene regulation compared to what was previously observed in UA159 grown with 0.5% fructose (33). As a control, similar treatment with 50 mM glucose for 30 min also upregulated 128 genes and down-regulated 216 genes (Table S3). Significantly, there was a larger overlap between the genes affected by fructose and methylglyoxal, totaling 176 genes (98 up and 62 down, 16 with opposite directions of change), compared to the 116-gene overlap between glucose and methylglyoxal (38 up and 48 down, 30 with opposite directions of change). Among the 176 genes shared by methylglyoxal and fructose, 61 overlapped the H_2_O_2_ regulon; among the 116 genes shared by methylglyoxal and glucose, only 24 overlapped with the H_2_O_2_ regulon (Table S5). Therefore, compared to glucose, significantly more genes were shared by treatment of fructose with that of methylglyoxal and H_2_O_2_. Comparative analysis identified a set of 44 genes that were similarly affected by fructose, methylglyoxal, and H_2_O_2_, excluding 17 genes impacted by all four treatments. Altogether, these 61 genes (Table S5 tab “Core genes”) encoded for some of the aforementioned functions such as thiol and metal homeostasis (*trxAB, gst, gor, sloA, dpr,* and *sufCDSUB*,), enzymes for ROS scavenging (*sodA, ahpCF,* and *tpx*) and restoration of redox balance (*nox*, ascorbate operon, *gapN*, and multiple *nrd* genes), glucan-binding protein *gbpC*, proteins implicated in general stress management including *clpE, clpL*, *uvrA, mutY,* SMU.503c (48), *exoA, lexA* (49), *pepP,* and *pepO* (50, 51), and transcription regulators such as *scnKR*, *perR,* and *msmR.* Again, nearly all 61 genes (except for 2) showed the same polarity of change under three treatments. Notably, many genes shared by glucose and fructose, e.g., *sloA, sodA, gor, tpx, gst, trxAB*, and *lexA*, showed significantly greater change due to the treatment by fructose than by glucose of the same amount.

Also included in this group of genes was the CRISPR1-Cas system (SMU.1402c to SMU.1406c) that was repressed by all four treatments, although the effect of H_2_O_2_ was slightly below the 2-fold threshold (Table S1). Conversely affected was the 10-gene CRISPR2-Cas cluster (SMU.1753c to SMU.1764c) that produced increased levels of transcripts under all four conditions, despite that the induction of several genes by methylglyoxal or glucose was below the threshold. An earlier study characterizing these two systems in *S. mutans* suggested that CRISPR-Cas overall was required for tolerance towards membrane, thermal, pH, and oxidative stressors (52). The differential regulation of these two systems in response to carbohydrates, methylglyoxal, and H_2_O_2_ warrants further exploration. It is also worth noting that the relative impact of fructose to both clusters was the highest among all four treatments.

#### Similarity between responses to fructose and glucose

Considering the effects of carbohydrates in increasing osmolarity and acid production, it was not surprising to find a substantial overlap between fructose- and glucose-responsive genes (total 165 genes, 80 genes when excluding methylglyoxal and H_2_O_2_; Supplemental Table S6), although many (near 50%) encoded hypothetical proteins. Interestingly, the operon required for malolactic fermentation (SMU.137-SMU.141) were most highly induced by either treatment, with the mRNA levels increasing by 12∼20-fold. Considering the inducibility of this system by acidic pH (53, 54), a rapid drop in pH due to glycolytic activities likely triggered this response. The agmatine deiminase (*aguA*) that has been shown to contribute to acid tolerance through production of ammonia (44) was similarly upregulated. Also induced by both carbohydrates were the majority of the genes/ORFs included in the genomic island TnSmu1 (55), from SMU.196c to SMU.215c, showing an increase in expression by 2∼4-fold. Other notably upregulated genes included the SMU.1602*/lguL* cluster, the zinc/cadmium-exporter *zccE* (56), the glucosyltransferase *gtfD*, the fructosyltransferase *ftf*, a putative F-1-P kinase *pfk* (17), the lactose operon (57), and the HPr kinase/phosphatase *hprK*. The fact that SMU.1602/*lguL* cluster was induced by methylglyoxal, fructose, and glucose suggested that endogenous production of methylglyoxal or related RES likely increased because of rapid influx of carbohydrates. Like before, *zccE* expression was >2-fold higher when treated with fructose compared to treatment by glucose. One cluster, SMU.1903c to SMU.1913c, along with the orphan response regulator *gcrR* (58), showed reduced expression under both treatments.

#### Genes affected only by fructose

A total of 168 genes were differentially affected by fructose alone (Supplemental Table S6 tab “Fru only”, Fig. S2A for enriched molecular functions). In addition to what have been discussed thus far, treatment with 50 mM fructose altered the expression of several pathways and additional genes required for carbohydrate metabolism. Namely, fructose upregulated the operons of *sppRA*, *fruRKI*, *levDEFG*, and the sorbitol PTS, however downregulated the nigerose PTS, the maltose PTS (*malT* or *ptsG*), *pfl,* the *msm* cluster (59), *galK*, *gtfC,* the glycogen operon (*glg*) (60), an α-glucan metabolic cluster (SMU.1564 to SMU.1571) (61), and importantly the fructose-bisphosphate aldolase (*fbaA*, down 2-fold) which is responsible for cleaving F-1,6-bP into dihydroxyacetone phosphate (DHAP) and glyceraldehyde 3-phosphate (GADP). Expression of three additional clusters was notably affected by fructose alone. First, SMU.1057 to SMU.1063, encoding SatCDE, Ffh, YlxM, and two ABC transporters, were upregulated by 2-to 3-fold. This cluster has been implicated in protein translocation and acid tolerance in *S. mutans* (62). Second, operon (SMU.1527 to SMU.1534) which encodes for the F_1_F_0_-ATPase complex were downregulated by 2-to 3-fold. It was reported that the F_1_F_0_-ATPase, whose main function in *S. mutans* is expelling excess proton to the environment, was upregulated in response to a shift from steady state at pH 7 to pH 5 as part of the acid tolerance response (ATR) (63–65).

Despite the likelihood of an acidic stress under the treatment of both glucose and fructose, there was no significant impact by glucose to this operon, and the treatment of fructose resulted in a lower rather than higher expression of all 8 genes encoding the F_1_F_0_-ATPase. Last, SMU.1734 to SMU.1745c, encoding for the pathway for fatty acid biosynthesis and similarly involved in ATR, showed 2- to 3-fold reduction in expression when treated with 50 mM fructose. While this effect was largely consistent with previous findings made in *S. mutans* under a glucose shock at 200 mM (65), here again 50 mM glucose failed to induce a similar response. Considering the importance of both pathways in maintaining membrane integrity and overall fitness, these findings could provide explanations to previously reported fructose-specific phenotypes such as autolysis and increased biofilm development (23, 30).

#### RT-qPCR analysis

Detailed transcriptional analyses were carried out to validate and further understand these findings, focusing on a list of genes relevant to bacterial fitness and pathogenicity. Considering the dynamic nature of gene regulation, mRNA abundance was compared in *S. mutans* UA159 treated with 50 mM fructose or 0.5 mM H_2_O_2_, each for 5 min and 30 min. Also included were the original treatments (30 min) of 5 mM methylglyoxal and 50 mM glucose, as well as two control conditions of growth in FMC-glucose and FMC-fructose. As shown in Table 1, mRNA levels of these 20 genes in the 4 treatments used in RNA-seq study confirmed their respective impact, including upregulation in functions such as radical scavenging, RES degradation, copper export, thiol homeostasis, and stress-responsive regulators; and downregulation of *mreC*, cellobiose PTS component *celB*, and mutacin gene *nlmA*. Thirty-min compared to 5-min treatment with H_2_O_2_ generally gave more substantial activation of most genes tested.

**Table 1.**
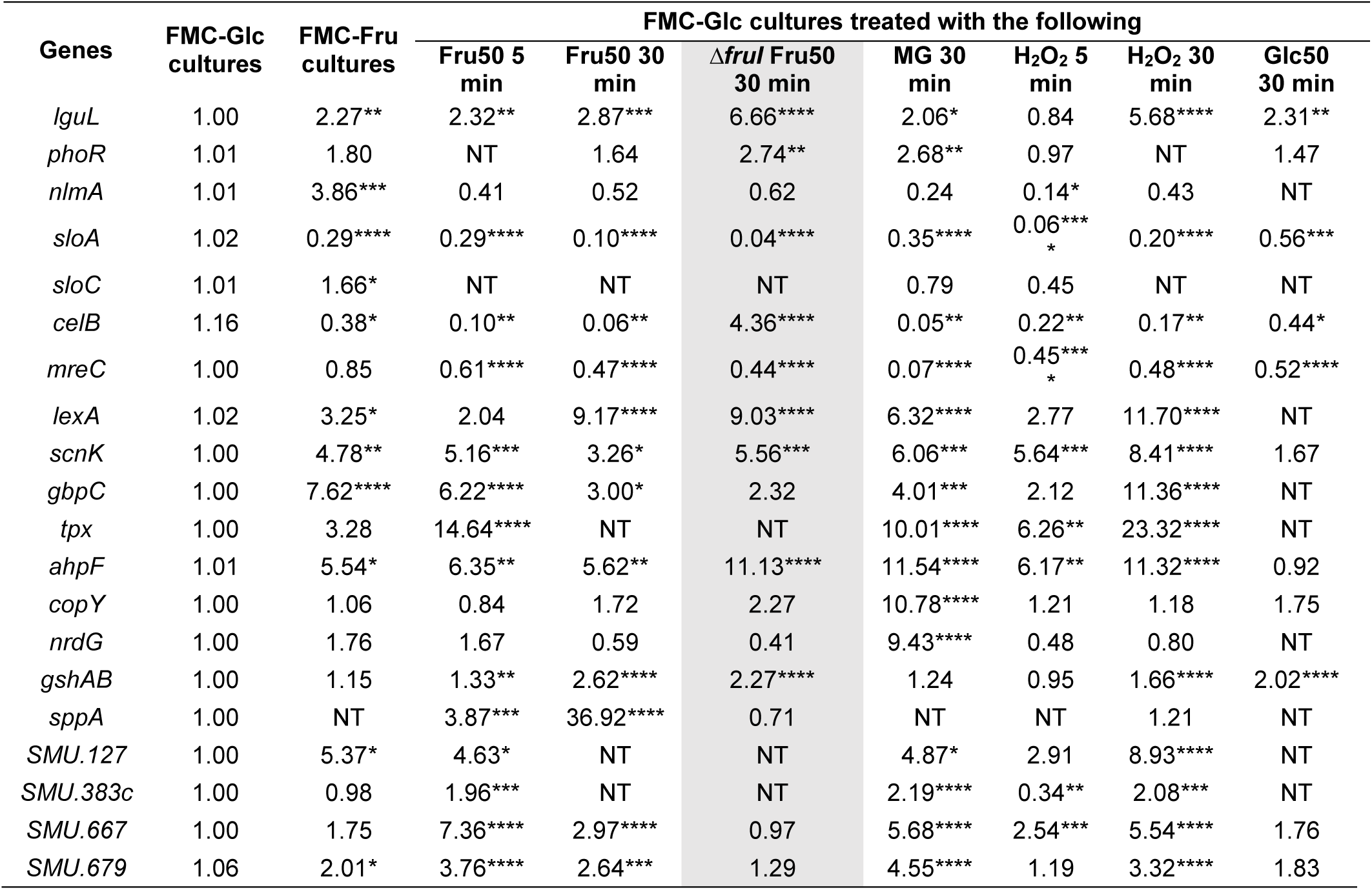
Relative expression of selected genes in *S. mutans* UA159 WT and Δ*fruI* (shaded) with specified treatments. After growing to mid-exponential phase (OD_600_ = 0.5) in FMC-Glc (20 mM glucose) at 37°C in 5% CO_2_, bacterial cultures were each treated with: Fru50, 50 mM fructose for 5 or 30 min; MG, 5 mM methylglyoxal for 30 min; H_2_O_2_, 0.5 mM hydrogen peroxide for 5 or 30 min; or Glc50, 50 mM glucose for 30 min. All values were relative to the untreated WT cultivated with FMC-Glc (second column). Exponential phase cultures of UA159 in FMC-Fru (20 mM fructose) were included as additional controls. Results are average from 3 biological replicates. Asterisks denote statistical significance assessed by One-Way ANOVA ((*, *P* <0.05; **, *P* <0.01; ***, *P* <0.001; ****, *P* <0.0001). NT, not tested.

Similarly, a time-dependent response to fructose was noted for genes *lguL*, *sloA*, *celB*, *lexA*, *gshAB,* and *sppA*, although several did show signs of reversal in impact, namely *scnK, gbpC, nrdG*, and SMU.667. To test the significance of F-1-P to gene regulation, a *fruI* null mutant was treated with 50 mM fructose for 30 min. To our surprise, only 4 out 16 tested genes showed a reversal in impact due to loss of FruI, suggesting that FruI-mediated effect plays only a partial role in the extensive transcriptomic reprogramming triggered by the fructose treatment.

### Fructose impacts metal homeostasis and Spx-dependent oxidative stress response

A likely effect of methylglyoxal on bacterial physiology was the reduction of intracellular GSH/GSSG ratio and potential impact on metal homeostasis. Previously it was demonstrated that a zinc/cadmium exporter ZccE was responsible for the high zinc tolerance of *S. mutans*, and its expression required a cognate regulator ZccR (56).

Deletion of either *zccE* or *zccR* resulted in a peroxide sensitivity under high-zinc conditions due to a disturbed zinc/manganese ratio which could be alleviated by manganese supplementation. When we tested the growth phenotype of a *zccR* null mutant in TV medium (66), which contains ∼19 μM of zinc according to ICP-OES analysis (56), the mutant nonetheless grew normally when supported by glucose as the sole carbohydrate source (Fig. 3A). When fructose was substituted for glucose however, Δ*zccR* showed a significant reduction compared to the WT UA159 in both growth rate and the final optical density (Fig. 3B). Like reported before, addition of 25 μM MnSO_4_ to the TV-fructose medium significantly alleviated the growth deficiency of Δ*zccR*. Importantly, after we introduced a *fruI* deletion into the *zccR* background, the double mutant grew at levels comparable to the WT UA159 on either glucose or fructose, without manganese supplementation (Fig. 3B). As *fruI* deletion was previously shown to reduce intracellular F-1-P levels (23), these findings strongly supported the theory that excess F-1-P levels negatively impacted metal homeostasis in *S. mutans*.

**Fig. 3.**
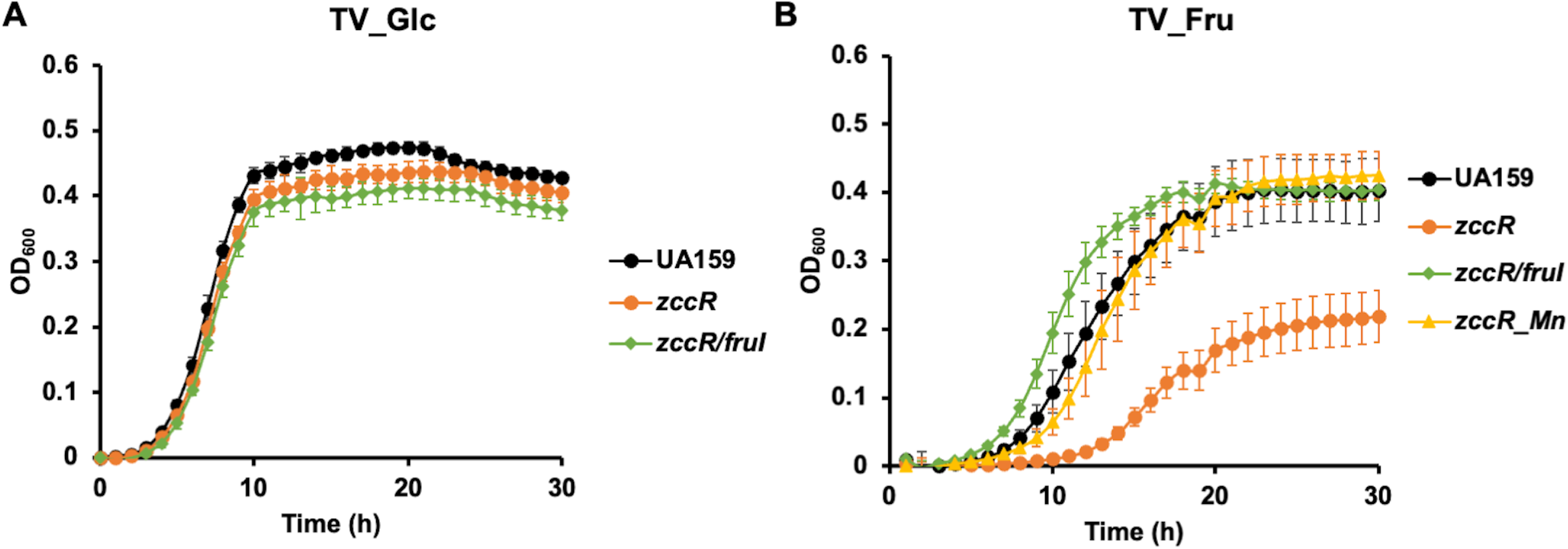
Fructose impacts metal homeostasis in Δ*zccR* deficient in zinc expulsion. *S. mutans* strains UA159 and its mutant derivatives were diluted from exponential phase cultures into TV supplemented with 20 mM glucose (A) or fructose (B). Mn^2+^ was added at 25 µM (MnSO_4_). Growth was monitored using a Synergy 2 plate reader maintained at 37°C. Results are average of three biological replicates, each tested in technical duplicates. Error bars denote standard deviations.

Since fructose-mediated transcriptome included a cohort of oxidative stress genes primarily regulated by SpxA1 (67), we repeated this growth assay by including a deletion mutant of *spxA1*. To our surprise, Δ*spxA1* grew at a faster rate in TV-fructose than in TV-glucose (Fig. 4A). Considering the relatively high zinc content in TV medium, we next switched to a chemically defined FMC medium that contained very little zinc but sufficient manganese (around 110 μM). In FMC medium, Δ*spxA1* grew relatively well on glucose but could not grow on fructose (Fig. 4B). When we introduced the same *fruI* deletion into the Δ*spxA1* background, the growth deficiency on fructose was greatly alleviated (Fig. 4C). At the same time, only modest benefit was observed on fructose when a *levD* deletion was introduced into Δ*spxA1.* Although the exact mechanisms remain to be clarified, the medium-specific fructose sensitivity of Δ*spxA1* added further support to the notion that F-1-P affects oxidative stress response and metal homeostasis by interacting with Spx-mediated regulation.

**Fig. 4.**
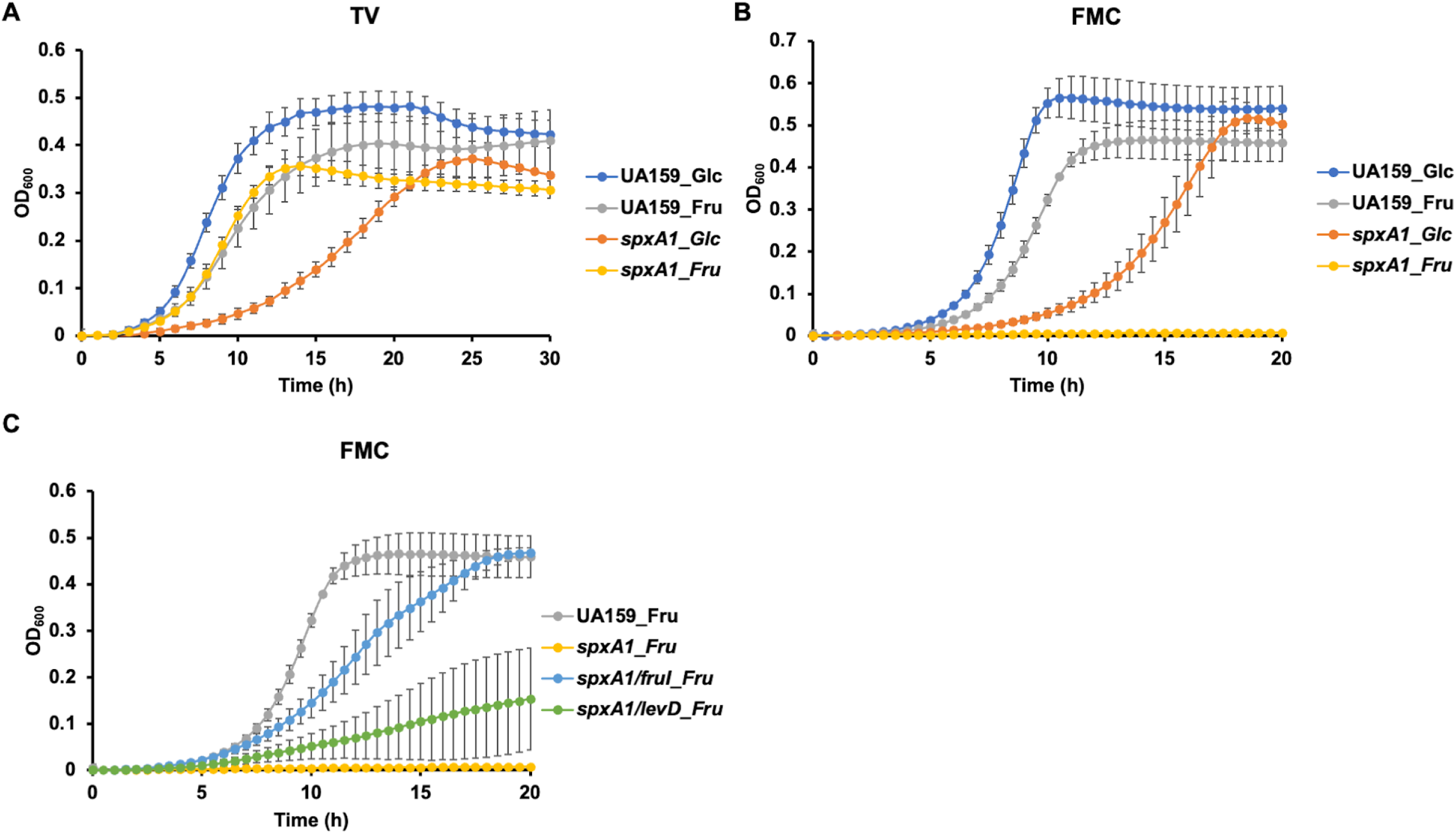
Fructose impacts growth of *spxA1* in a medium-specific manner. *S. mutans* strains UA159 and various mutant derivatives were each diluted from exponential phase cultures into TV (A) or FMC (B, C) supplemented with 20 mM glucose (Glc) or fructose (Fru). Growth was monitored using a Synergy 2 plate reader maintained at 37°C. Results are average of three biological replicates, each tested in technical duplicates. Error bars denote standard deviations.

To further test this hypothesis, a P*sodA::gfp* promoter-reporter fusion was established in the WT UA159 background to assess the response of Spx-mediated regulation to the presence of fructose; the superoxide dismutase (*sodA*) requires Spx proteins for optimal expression (36). As shown in Fig. 5 and Fig. S3, the *sodA* promoter was activated by the presence of fructose in a dose-dependent manner, from levels as low as 0.5 mM, yet it only produced a modest and slightly delayed response to the presence of 0.5 mM of H_2_O_2_. While 0.5 mM H_2_O_2_ significantly impeded the growth of *S. mutans* [Fig. S4 and (68)], as much as 10 mM fructose showed little to now effect. To ascertain the nature of fructose-derived signal in this effect, *fruI* and *levD* deletions were each introduced into UA159 containing the P*sodA::gfp* fusion. Consistent with above observations, deletion of *fruI* but not *levD* resulted in the loss of fructose-dependent activation of the *sodA* promoter activity. As expected, deletion of *spxA1* also abolished the expression of *sodA* promoter (36). These results not only confirmed the role of F-1-P in influencing Spx regulation, but given the magnitude of impact, suggested that fructose may serve as a primary signal for bacterial response to environmental changes.

**Fig. 5.**
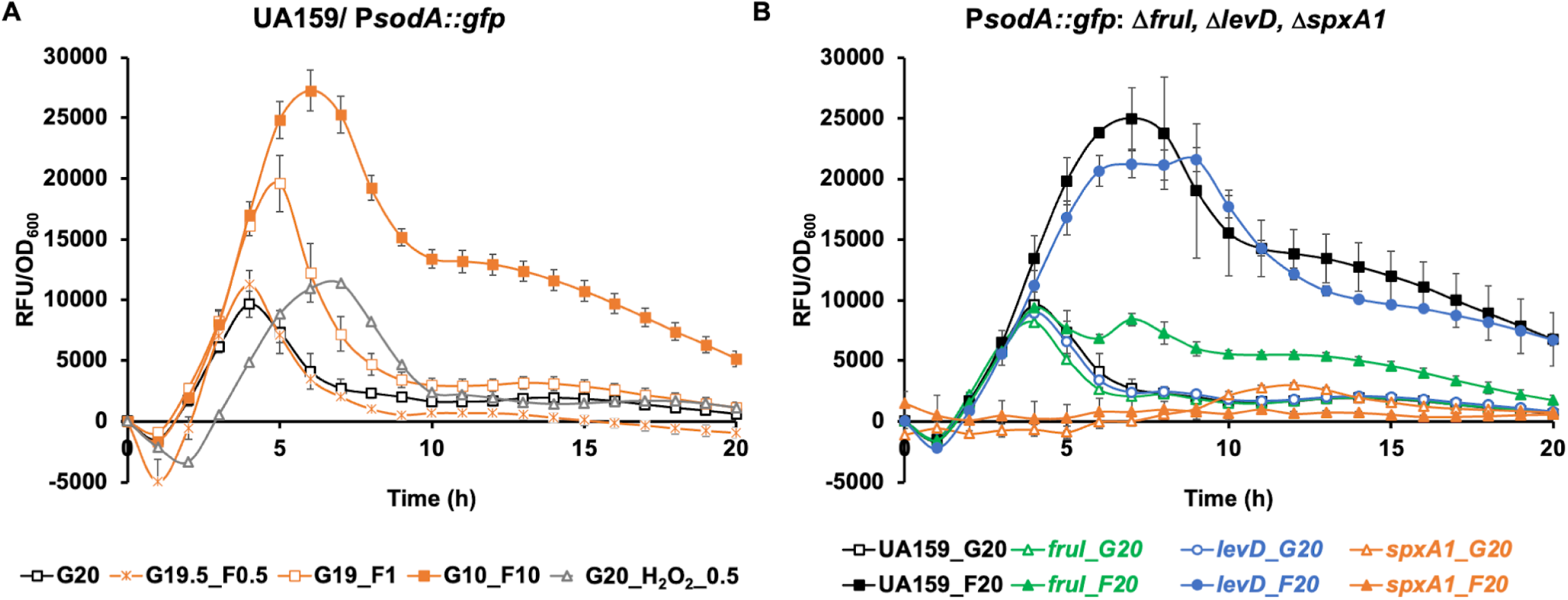
Induction of the *sodA* promoter by fructose. (A) Cultures of UA159 harboring a P*sodA::gfp* fusion were diluted into FMC containing specified concentrations (in mM) of glucose (G), fructose (F), or H_2_O_2_. The cultures were monitored in a Synergy 2 plate reader for green fluorescence and optical densities (OD_600_) for 20 hours. (B) The same experiment was performed on various mutants derived from UA159/P*sodA::gfp,* using FMC containing 20 mM glucose (G20) or fructose (F20). The relative fluorescence units (RFU) of each culture were recorded as a measure of *sodA* promoter activity, subtracted with RFUs of a control strain (the same genetic background but without the *gfp* fusion) cultured under the same condition, and normalized against corresponding OD_600_ values of the bacterial cultures. Results are the average from three biological replicates, conducted in technical duplicates. Error bars denote standard deviations.

F-1-P is the likely allosteric inducer for activating the *fruRKI* operon. To assess the levels of fructose needed to induce the expression, we utilized a strain containing a promoter-reporter fusion P*fruR::cat* (20). Bacteria were cultured in glucose-based FMC medium and then exposed to different amounts of fructose for 30 min. CAT activities from UA159/P*fruR::cat* suggested that at levels as low as 20 μM, fructose can significantly induce the expression of the *fruRKI* operon (Table S7).

### Fructose contributes to fitness during starvation, acidic pH, and competition with peroxigenic streptococci

Our previous studies have suggested that fructose, when used in excess under laboratory conditions, may induce autolysis and cell death. Paradoxically, considering the activation of much of the stress response core by fructose, we could anticipate improved persistence under stressful conditions. To test this hypothesis, we subjected *S. mutans* to a series of physiological tests by exposing it to fructose. First, strain UA159 was cultivated in FMC with 20 mM glucose or fructose, and their CFU was monitored while these cultures were continuously incubated without medium refreshment for 11 days. While fructose produced lower CFU than glucose at day 1 of the incubation as reported previously (23), it maintained the CFU of UA159 by at least 1 log higher than on glucose throughout the rest of the experiment (Fig. 6A). When 1 mM fructose was used together with 19 mM glucose, the fructose-associated effects were reduced and limited to the first two days of the incubation. We then repeated this experiment using strain Δ*fruI* (Fig. 6B). Echoing the findings in RT-qPCR analysis, Δ*fruI* showed significantly higher persistence than the wild-type parent, especially when fructose was used.

**Fig. 6.**
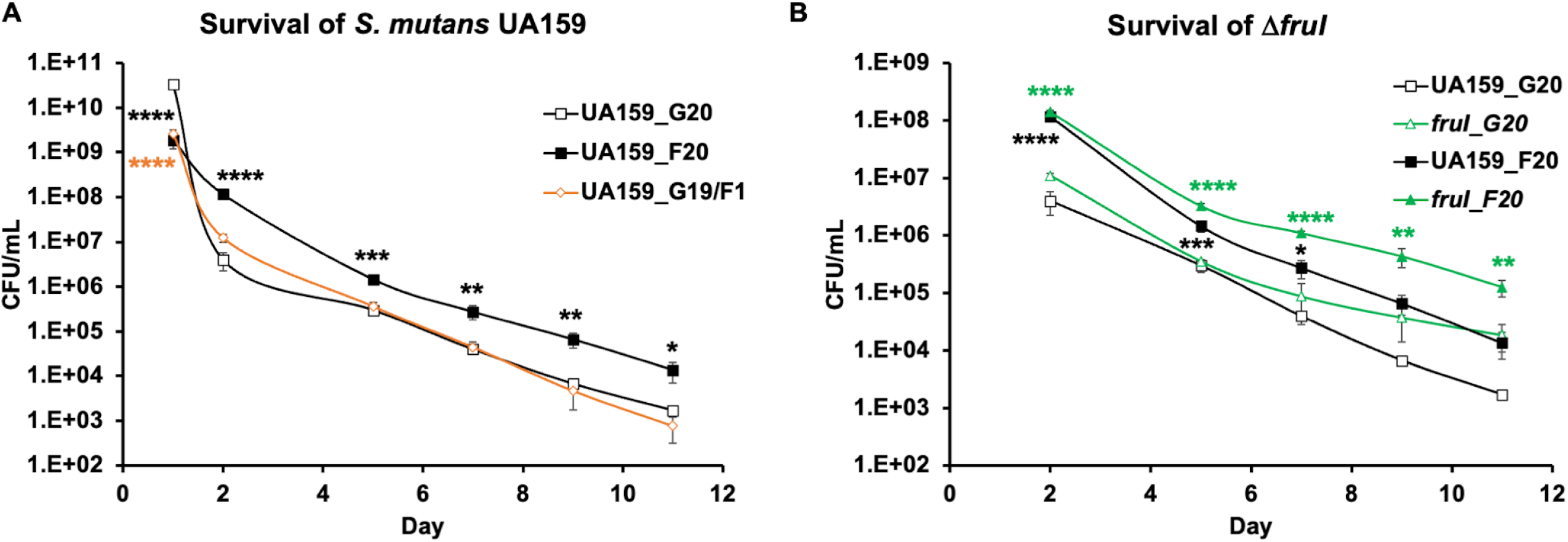
Fructose promotes long-term survival of *S. mutans*. Cultures of *S. mutans* UA159 (A, B) or Δ*fruI* (B) were diluted into FMC containing 20 mM of glucose (G20) or fructose (F20), or a combination of 19 mM glucose and 1 mM fructose (G19/F1), followed by incubation without medium refreshment for 11 days. At specified time points, an aliquot of culture was removed for CFU enumeration. Each strain was represented by 3 independent cultures, with results presenting average and standard deviation. Statistical significance was assessed using (A) One-Way ANOVA followed with Dunnett’s comparisons against G20 samples, and (B) Two-Way ANOVA followed by Tukey’s multiple comparisons against G20 samples (*, *P*<0.05; **, *P*<0.01; ***, *P*<0.001; ****, *P*<0.0001).

As a further test of acid tolerance under the fructose condition, we cultured UA159 for 20 h in a TV medium (FMC contains 10 mM phosphate buffer thus not ideal) supported with different amounts of glucose or fructose and measured the resting pH of these cultures. The results (Table 2) showed comparable growth yield but significantly lower resting pH in TV-fructose cultures than in TV-glucose cultures. For comparison, we conducted the same experiment on a health-associated commensal, *S. sanguinis* (SSA) strain SK36. To our surprise, SK36 failed to produce a substantial growth on fructose, especially for higher concentrations (100 or 200 mM). Similar findings were made when this experiment was repeated using another *S. sanguinis* isolate BCC23 (69). Further tests on a number of low-passage, clinical isolates of additional commensal streptococci (70), encompassing the species of *S. gordonii, S. mitis, S. oralis, S. dentisani, S. intermedius, S. sp.* A12*, S. parasanguinis,* and *S. cristatus*, showed similar sensitivity in most but not all species (e.g., *S. intermedius*) tested (Table 2). On the contrary, tests on 6 additional WT *S. mutans* strains (Table S8) indicated that most of them were able to grow comparably well on fructose, and lowered the environmental pH to similar levels to that achieved on glucose. However, one isolate, OMZ175 of serotype *f* known for its collagen-binding and invasive activities (71), did show a modest sensitivity to fructose by having lower final OD_600_ and notably higher resting pH on fructose than glucose. We then performed a pH drop assay to further test the notion that fructose could lower the culture pH better than glucose. When a suspension of UA159 cells from an exponential phase, fructose-grown culture was fed 50 mM of fructose without buffering, it lowered the environmental pH at a notably faster rate, which ended at a significantly lower point, than a glucose-grown culture of UA159 did in 50 mM glucose (Fig. 7A and Fig. S5). It appeared that *S. mutans* UA159 catabolized fructose more rapidly than glucose and was more acid-tolerant under fructose conditions. We also performed this assay using BHI-grown bacteria for pH drops with glucose or fructose, which showed a similar, albeit smaller difference between these two sugars. We conclude that while most *S. mutans* strains tested by us appear well adapted to fermenting fructose, *S. sanguinis* and many commensal streptococci may be less capable of tolerating high levels of fructose.

**Table 2.** pH and final OD_600_ of 20-h TV cultures.

| Strains | Sugars | Resting pH |  |  | Final OD<sub>600</sub> |  |  |
| --- | --- | --- | --- | --- | --- | --- | --- |
|  |  | 20 mM | 100 mM | 200 mM | 20 mM | 100 mM | 200 mM |
| SMU-UA159 | Glucose | 4.92 | 4.93 | 4.93 | 0.83 | 0.72 | 0.57 |
|  | Fructose | 4.76**** | 4.82** | 4.81*** | 0.82 | 0.73 | 0.64**** |
| SSA_SK36 | Glucose | 6.5 | 6.15 | 5.70 | 0.34 | 0.41 | 0.45 |
|  | Fructose | 6.55 | ND | ND | 0.27*** | 0.03**** | 0.00**** |
| SSA_BCC23 | Glucose | 6.09 | 6.01 | 5.93 | 0.43 | 0.37 | 0.31 |
|  | Fructose | 5.76*** | ND | ND | 0.49* | 0.20**** | 0.02**** |
| SDE_BCC10 | Glucose | 5.48 | 5.08 | 5.49 | 0.63 | 0.69 | 0.43 |
|  | Fructose | ND | ND | ND | 0.28**** | 0.14**** | 0.00**** |
| SOR_BCC11 | Glucose | 5.72 | 5.65 | 5.72 | 0.56 | 0.52 | 0.4 |
|  | Fructose | 5.09**** | 5.56 | ND | 0.66**** | 0.55 | 0.01**** |
| SPA_BCC15 | Glucose | 6.53 | 6.45 | 6.38 | 0.12 | 0.11 | 0.09 |
|  | Fructose | ND | ND | ND | 0.02*** | 0.00*** | 0.00** |
| SMI_BCC2 | Glucose | 4.75 | 4.73 | 4.77 | 0.74 | 0.65 | 0.56 |
|  | Fructose | 4.87** | 4.87*** | 4.87** | 0.61**** | 0.54**** | 0.47**** |
| SGO_BCC9 | Glucose | 4.95 | 4.90 | 4.91 | 0.81 | 0.66 | 0.54 |
|  | Fructose | 4.87 | 5.03 | 5.97**** | 0.70**** | 0.68 | 0.46*** |
| A12_BCC21 | Glucose | 5.62 | 5.37 | 5.48 | 0.65 | 0.56 | 0.46 |
|  | Fructose | 5.16*** | 5.36 | ND | 0.64 | 0.54 | 0.28**** |
| SCR_BCC41 | Glucose | 6.20 | 5.93 | 5.85 | 0.5 | 0.47 | 0.42 |
|  | Fructose | 5.69**** | 5.95 | ND | 0.51 | 0.41 | 0.10**** |
| SIN_BCA19 | Glucose | 5.28 | 5.12 | 5.18 | 0.66 | 0.58 | 0.48 |
|  | Fructose | 5.11*** | 5.01* | 5.14 | 0.61**** | 0.55** | 0.48 |

Results show the averages of four independent cultures. Statistical significance was determined using Two-Way ANOVA followed by Bonferroni's test for comparisons with glucose cultures of the same concentration (\*, $P < 0.05$ ; \*\*, $P < 0.01$ ; \*\*\*, $P < 0.001$ ; \*\*\*\*, $P < 0.0001$ ). ND, not determined due to lack of substantial growth (in red). SMU, *S. mutans*; SSA, *S. sanguinis*; SDE, *S. dentisani*; SOR, *S. oralis*; SPA, *S. parasanguinis*; SMI, *S. mitis*; SGO, *S. gordonii*; A12, *S. sp. A12*; SCR, *S. cristatus*; SIN, *S. intermedius* (70, 100).

**Fig. 7.**
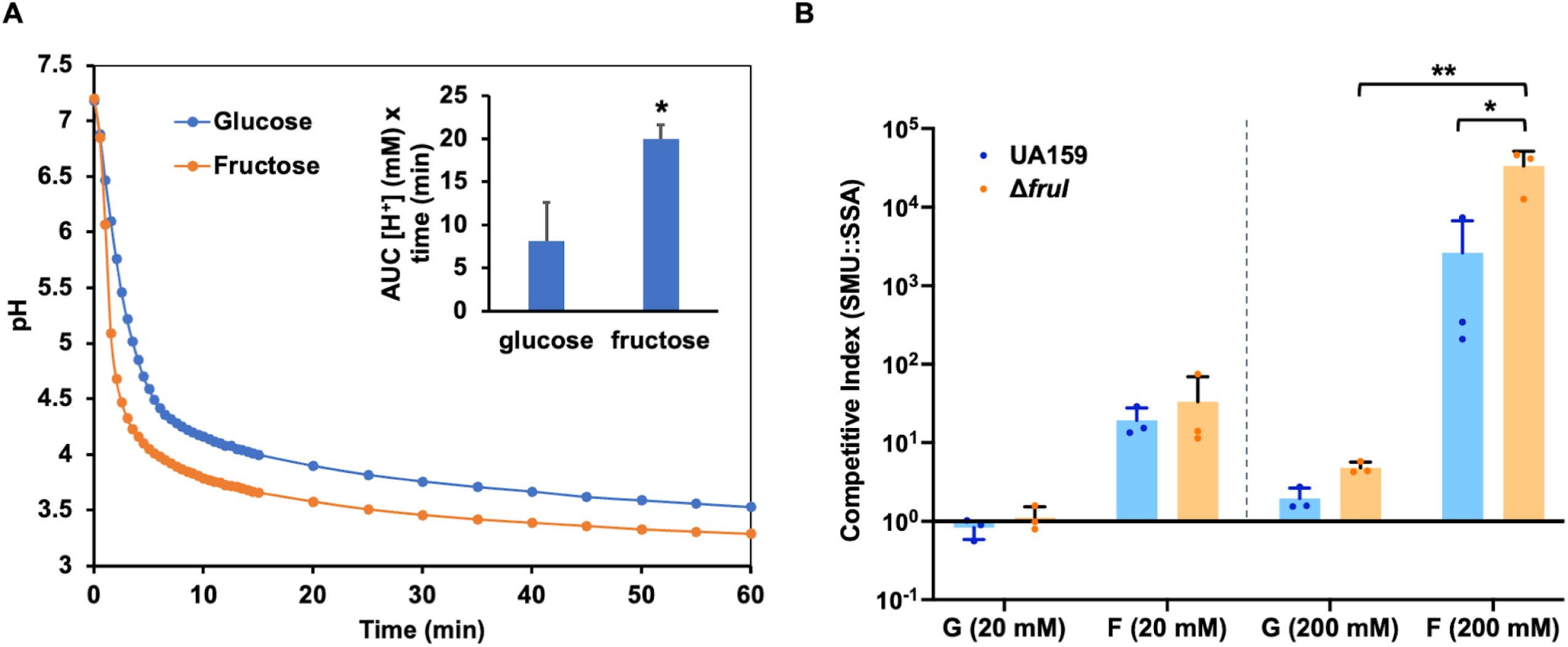
Fructose enhances acidogenicity (A) and competitiveness (B) of *S. mutans*. (A) pH drop assay. UA159 was cultured in TV with 20 mM glucose or fructose to exponential phase, harvested and resuspended in a solution of 50 mM KCl and 1 mM MgCl_2_, with OD_600_ adjusted to 5.0. The pH was adjusted to 7.2 slowly with 0.1 N KOH to consume intracellular carbohydrate storage. Immediately after addition of 50 mM glucose or fructose, same as the sugar used in the TV medium, the pH of the suspension was monitored continuously on a stirring plate for the next 60 min. Each experiment was repeated three times (Fig. S5), with representative results shown here. After conversion into proton concentrations, area under the curve (AUC, see inset) of each sample was calculated and used to assess statistical significance using Student’s *t*-test (*, *P* < 0.05). (B) Cultures of *S. mutans* wild type UA159 or Δ*fruI* were mixed at 1:1 ratio with a differentially marked *S. sanguinis* strain, then diluted at 1:100 into FMC containing glucose (G) or fructose (F), each used at 20 mM or 200 mM. After 24 h of incubation, the cultures were sonicated and used for CFU enumeration. CFUs from both the start and end of the incubation (Fig. S6) were used to calculate competitive indices. Results were the average of three independent repeats, with error bars denoting standard deviations. Statistical significance was assessed using Two-Way ANOVA followed by Tukey’s multiple comparisons (*, *P*<0.05; **, *P*<0.01).

Next, *S. mutans* was subjected to a competition assay in a mixed-species planktonic culture with the peroxigenic commensal *S. sanguinis*. Cultures from the exponential phase were mixed at approximately 1:1 ratio between two species and diluted into the FMC medium supplemented with 20 mM of glucose or fructose. After an overnight of incubation, CFUs for each species were enumerated by plating on selective agar plates (Fig. S6 includes CFU from mixed-species cultures and single-species controls). The resultant competitive indices (Fig. 7B) indicated a greater competitiveness for *S. mutans* UA159 in the presence of fructose than glucose, especially when used at higher concentrations. It is perhaps worth noting that this competitiveness of UA159 favored by fructose was at least partly attributable to the fructose sensitivity of its competitor, SK36, a phenotype that could be species- and even strain-specific (Table 2). Interestingly, deleting *spxB* in SK36 did not significantly change the competitive indices (Fig. S6E, 20 mM carbohydrates) whereas deleting *fruI* in UA159 further enhanced the competitiveness of *S. mutans* against *S. sanguinis* SK36 when supported with fructose. Since Δ*fruI* has a defect in fructose transport and grows more slowly than the WT on fructose (17), these and earlier results (Fig. 6B) suggested that a certain optimal level of F-1-P could benefit *S. mutans* when competing against commensal species capable of producing inhibitory levels of H_2_O_2_.

## Discussion

It is frequently observed in stress biology that different stressors result in overlapping responses in bacteria (72). As an oral bacterium, *S. mutans* is frequently subjected to environmental insults that include acidic pH, oxygen and H_2_O_2_, hyperosmolarity, thermal fluctuation, metal ions, and nutrient starvation (73). In this study, we identified methylglyoxal and fructose as two potential stress signals capable of inducing streptococcal stress responses (Fig. 8). A stress core comprised of 61 genes were shared by the regulons of fructose, methylglyoxal, and H_2_O_2_. This commonality in gene regulation can be attributed to the overlapping effects exerted by these treatments. For example, exposure to concentrated sugars likely leads to acidic stress, hyperosmolarity, and increased RES production. Treatments by both RES and ROS are expected to perturb intracellular redox pools that could in turn influence bacterial metal and thiol homeostasis. Exposure to extreme pH, RES, and ROS can all lead to denaturation of amino acids, proteins, lipids, and nucleic acids. Each of these outcomes likely activates a central regulatory system to restore essential nutrients or repair damage. Not surprisingly, 40 out of the 61 core stress genes (Table S5) have been previously identified in *S. mutans* by an acid adaptation study in response to either a glucose shock (200 mM) or a shift from neutral to acidic pH (65). Conversely, only 17 out of the 61 stress-related genes were affected by our treatment of 50 mM glucose. The discrepancy between our study and earlier findings by others (65) is likely due to the difference in sugar content and culture conditions, as 200 mM glucose and continuous culturing were used before as opposed to 50 mM glucose and batch cultures herein. Corroborating the transcriptomic overlap between fructose and acid tolerance, phenotypic analyses suggested a superior ability for *S. mutans*, particularly certain strains such as UA159, GS-5 and ST1 (Table S8), to lower environmental pH when fructose was used as the supporting carbohydrate, especially at higher concentrations. This ability to tolerate fructose appears to be limited or absent in many commensal streptococcal species we have tested so far, with potential species and strain specificity, which may have important implications to microbial ecology in oral biofilms. Furthermore, several pathways triggered by hyperosmolarity in previously studies (74–76), including genes *lguL, sodA, nox*, the *opcA* cluster (SMU.2116 to SMU.2119), the *opu/ffh/sat* cluster, and potassium uptake mechanism *trkAB*, were also identified as part of the fructose/methylglyoxal/H_2_O_2_ response core.

**Fig. 8.**
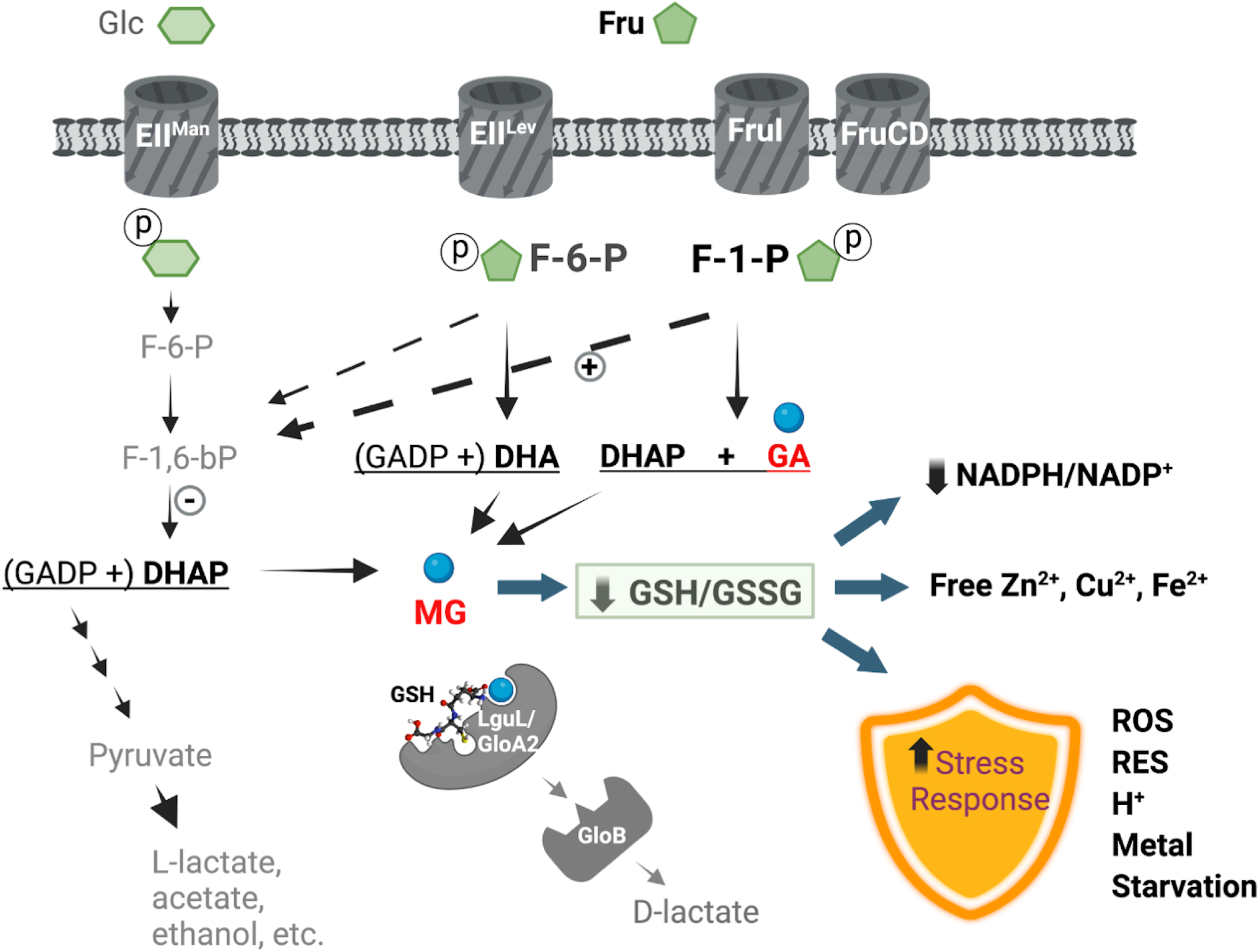
Fructose metabolism triggers stress response in streptococci. Fructose is internalized by oral streptococci primarily through a F-1-P-generating PTS transporter, FruI, but also through two additional PTS permeases, including EII^Lev^ (LevDEFG) that generates F-6-P. Compared to the glucose pathway, fructose is likely catabolized more rapidly via glycolysis, during which RES such as methylglyoxal is accumulated and impacts cytoplasmic redox balance represented by the ratios of GSH/GSSG and NADPH/NADP^+^. Increased F-1-P kinase activities (denoted by a plus sign) and reduced expression of F-1,6-bP aldolase (by a minus sign) may further contribute to RES production through impeded F-6-P phosphorylation and increased cleavage of F-6-P into DHA. Also affected are the pools of intracellular metal ions. Sensing these and other not-yet-characterized physiological changes and specific metabolites, the bacterium reprograms a suite of stress responsive genes. The outcomes of exposure to fructose include not only elevated activities in processing RES, through the functions of LguL/GloB and related protein GloA2, but importantly enhanced tolerance to multiple environmental stressors including ROS, RES, acidic pH, toxic metals, and nutrient depletion.

Theories established in mostly Gram-negative bacteria suggest that methylglyoxal bypass, a function that converts DHAP into pyruvate via the activities of a methylglyoxal synthase (*mgsA*), two glyoxalases (*gloAB*), and a D-lactate dehydrogenase (D-*ldh*), serves an important function in modulating glycolysis under sugar excess and phosphate limitation (32). Genes *mgsA* and D-*ldh* are absent in most lactic acid bacteria (LAB) genomes (30). However, methylglyoxal and related RES can be produced from DHAP spontaneously or from a variety of other mechanisms (77). The conservation of glyoxalases in streptococci (78) and the transcriptomic overlap among fructose, glucose, and methylglyoxal support the notion that methylglyoxal metabolism is an integral part of bacterial physiology, and the sources of methylglyoxal could be both endogenous and exogenous. This conclusion is consistent with an emerging theory in bacterial pathophysiology termed the aldehyde hypothesis (79), where a group of reactive metabolic intermediates (RES) from the host and pathogens may serve as potential antimicrobial effectors. This theory has supports from studies on LAB pathogens that engage host immunity, including *S. pyogenes* and *S. agalactiae*, which required glyoxalases for fitness and virulence (31, 78). For oral streptococci, the effects of fructose on methylglyoxal metabolism could involve several mechanisms. First, multiple RES compounds can be generated during fructose metabolism, given the presence of three phosphorylation products, F-1-P, F-6-P, and F-1,6-bP. Glucose metabolism generates only F-6-P and F-1,6-bP. DHAP is a direct product of cleavage of F-1,6-bP; cleavage of F-1-P by an aldolase produces glyceraldehyde (GA) and DHAP; and similarly, cleavage of F-6-P yields G3P and dihydroxyacetone (DHA). These metabolites are either themselves RES, e.g., GA, or in the case of DHAP and DHA, can generate methylglyoxal due to a reaction that is spontaneous or enzymatic in nature (MgsA). Second, fructose degradation may be regulated differently than glucose degradation, which could affect the accumulation or degradation of metabolic intermediates that are precursors of RES. Rapid phosphorylation of fructose by fructokinase and the resultant drawdown of ATP is a major factor in fructose-dependent metabolic impact in mammalian cells (16). Although in bacteria, carbohydrates are phosphorylated by PTS at the expense of PEP instead of ATP, here we showed a notably faster rate of glycolysis by *S. mutans* UA159 degrading fructose than glucose. As a result, fructose influx may alter the metabolism of F-6-P and F-1,6-bP, impacting RES production and gene regulation even in Δ*fruI* background. For example, accumulation of F-6-P and its degradation into DHA (instead of phosphorylation into F-1,6-bP) could presumably be amplified during high-fructose treatment, especially if F-1,6-bP cleavage is impeded. In support of these reasonings, fructose transcriptome reported downregulation of the F-1,6-bP aldolase *fbaA* (by 2-fold), and many common genes shared by fructose and glucose treatments produced a greater amplitude of change under the fructose condition. As part of the effort to understand the genetic mechanisms of fructose-specific biology, we have identified a novel 5-gene cluster, well conserved in *S. mutans* and most streptococci, which encodes a putative F-6-P aldolase and a glycerol dehydrogenase (80, 81). Products of these genes could contribute to metabolism of F-6-P and DHA, as suggested by a study in *Listeria innocua* (82); and their overexpression was recently implicated in the hypervirulent phenotypes of a dominant GAS variant in England (83). Moreover, most Gram-positive bacteria maintain a novel LguL paralog, GloA2, whose deletion in *S. mutans* resulted in a fructose-specific phenotype that resembled methylglyoxal exposure (30). Highly pertinent to our research, an earlier animal study demonstrated that high fructose corn syrup (HFCS, 45% glucose and 55% fructose), despite producing no glucans, can induce dental caries in a rat model as effectively as sucrose (84). Likewise, *in vitro* studies by others reported that HFCS induced greater demineralization of tooth enamel compared to a similar treatment by sucrose (85).

In addition to influencing RES metabolism, F-1-P homeostasis appears to be an essential physiological signal in most streptococci. The *fruRKI/fruRBA* operon required for catabolizing fructose via the F-1-P pathway is highly conserved in most Gram-positive bacteria. Several oral streptococci also harbor the *sppRA* operon for moderating the F-1-P levels (23). We previously showed that a *fruK* deficient strain accumulated significant levels of F-1-P even at the absence of fructose and was highly defective in growth on fructose, suggesting an endogenous source of F-1-P (e.g., dephosphorylation of F-1,6,-bP) and the need to keep its level within an optimal range (23). Notably, the *sppRA* promoter required higher levels of fructose (25 to 50 mM) for induction (23), suggesting that *sppRA* are expressed only when F-1-P is above a certain threshold to avoid futile cycling between F-1-P and fructose. Conversely, we have determined herein that 0.5 mM fructose is capable of activating Spx-mediated regulation and as little as 20 μM of fructose efficiently induced the expression of the *fruRKI* operon. We also observed in *S. sanguinis* the induction of *fruRBA* expression by similar levels of fructose or 25% human sera (Table S7). It is worth noting that for diabetic or hyperglycemic individuals, blood fructose levels are often significantly elevated as a function of the polyol pathway (16), capable of activating the *fruRKI/fruRBA* pathway (86). These findings allow us to posit that fructose regulates streptococcal function at both high (mM) and low (μM) levels, with the former contributing to fitness during the feast-and-famine cycle of the oral cavity and the latter in human circulatory system. While we only showed that fructose can benefit *S. mutans* at mM concentrations, we have good reasons to speculate that a similar benefit applies under μM levels of fructose. Several studies have suggested that *fruRKI/fruRBA* genes are required for streptococcal virulence during extraoral infections (19, 22, 83), where only sub-mM fructose is available, although the mechanisms remain uncharacterized. We further posit that fructose may trigger a surveillance mechanism against host-derived ROS and RES molecules, which primes bacterial stress response and additional fructose-specific pathways (Fig 8). Aside from FruR, SppR, LevQRST, and LacR that were previously shown to be responsive to fructose, herein we identified several transcription regulators whose expression appeared uniquely responsive to fructose treatment, including *spxA2, glnR, hdrR,* and a two-component system *spaRK*. Targeting these regulators and F-1-P-specific metabolism, i.e., GloA2, could help us unravel the molecular mechanisms required for cells to optimize stress management in response to fructose. With many oral streptococci existing as dual-niche organisms and the widespread use of fructose in food, this research has implications in both oral microbial homeostasis and systemic health.

Lastly, it is worth reiterating that a fructose-PTS mutant (*fruI*) of *S. mutans* displayed reduced colonization and cariogenicity in an earlier rat study under high-sucrose condition (24). This study revealed an important connection between fructose and sucrose in *S. mutans* physiology, as sucrose metabolism releases significant quantities of fructose both intracellularly and extracellularly, the latter of which can be reinternalized by the PTS (33, 34, 87). While it is tempting to speculate that fructose released from sucrose might contribute to virulence of *S. mutans* and influence the persistence of certain commensal streptococci, more in-depth investigation is needed to fully understand the role of fructose in caries development and biofilm dysbiosis.

### Concluding remarks

By establishing a stress regulon shared by fructose, methylglyoxal, and H_2_O_2_, our study revealed a unique identity of fructose in streptococcal physiology as a fermentable substrate, a stressor, and potentially an important environmental signal. Gene regulations associated with an F-1-P homeostasis contributed to bacterial fitness under stressful conditions, especially for the cariogenic pathobiont *S. mutans*. The overlapping nature of stress response in general also suggests that fructose metabolism may affect the overall resilience against other stressors such as acidic pH, ROS, RES, toxic metals, starvation, and hyperosmolarity; conditions to be expected in both oral cavity and the blood stream (Fig. 8). Considering the widespread conservation of some of these genetic features, most of the fructose-mediated effects discussed here are likely applicable to other lactic acid bacteria which have high relevance in both oral and systemic health.

## Materials and Methods

### Bacterial strains and culture conditions

Brain heart infusion (BHI) medium (Difco Laboratories, Detroit, MI) was used to maintain *S. mutans* and other streptococcal strains (Table 3). Liquid BHI was used to prepare for starter cultures for most of the experiments. Antibiotics were used, only when necessary, at the following concentrations: kanamycin (Km) 1 mg/mL, erythromycin (Em) 10 µg/mL, and spectinomycin (Sp) 1 mg/mL. For cultures requiring specific carbohydrates, a semi-defined Tryptone-Vitamin (TV) medium (66) or a synthetic medium FMC (88) was used, each supplemented with specified carbohydrates. Selection between TV and FMC was made based on the requirements of each experiment. Specifically, FMC was selected as the base medium when certain carbohydrates were added at very low levels, 1 mM or lower, to avoid potential contaminations from ingredients of TV. Similarly, FMC was used in promoter reporter study to reduce background fluorescence. TV and FMC also contain significantly different buffering capacity and levels of nutritional and toxic metal ions. Other than growth curves, all cultures were incubated without agitation at 37°C in an aerobic atmosphere supplemented with 5% of CO_2_.

**Table 3.** Bacterial strains used in this study (excluding clinical isolates used in Table 2).

| Strains | Relevant characteristics<sup>a</sup> | Source or reference |
| --- | --- | --- |
| *S. mutans* UA159 | *S. mutans* wild type, *perR*<sup>+</sup> | ATCC 700610 |
| MMZ1654 | UA159 *frul::Km* | From UA159 |
| MMZ2120 | UA159 *frul::Em* | From UA159 |
| $\Delta zccR$ | UA159 *zccR::Sp* | (56) |
| MMZ2156 | UA159 *zccR::Sp frul::Km* | From $\Delta zccR$ |
| $\Delta spxA1$ | UA159 *spxA1::Sp* | (36) |
| MMZ2161 | UA159 *spxA1::Sp frul::Em* | From $\Delta spxA1$ |
| MMZ2162 | UA159 *spxA1::Sp levD::Em* | From $\Delta spxA1$ |
| $\Delta spxA2$ | UA159 *spxA2::Em* | (36) |
| MMZ2158 | UA159 *sodA::gfp* fusion (*Km*) | From UA159 |
| MMZ2160 | UA159 *spxA1::Sp sodA::gfp* fusion | From $\Delta spxA1$ |
| MMZ2159 | UA159 *frul::Em sodA::gfp* fusion | From MMZ2120 |
| MMZ2173 | UA159 *levD::Em sodA::gfp* fusion | From MMZ2158 |
| MMZ799 | UA159 *fruR::cat* fusion (*Km*) | (20) |
| UA159-*Km* | UA159 *gtfA::Km* | (96) |
| *S. sanguinis* SK36 | *S. sanguinis* wild type | Kitten laboratory |
| MMZ1945 | SK36 *gtfP::Em* | (97) |

<sup>a</sup> *Km*, *Em*, and *Sp*: resistance against kanamycin, erythromycin, and spectinomycin, respectively.

For growth curves, bacterial cultures were diluted 100-fold into a TV or FMC medium containing specified carbohydrates, overlaid with 70 µL mineral oil, and maintained at 37°C in a Bioscreen C lab system (Labsystems Oy, Helsinki, Finland) that recorded the optical density of the cultures at 600 nm (OD_600_) every hour. A brief lateral shaking was included before each reading to disperse aggregates. To monitor the expression levels of the *sodA* promoter over time using a P*sodA::gfp* reporter fusion, bacterial cultures from the exponential phase were similarly diluted into an FMC medium containing specified amounts of carbohydrates and H_2_O_2_ and overlaid with mineral oil. For 20 h, the OD_600_ and the relative fluorescence units (RFU) of the cultures (excitation 485/20 nm, emission 528/20 nm) were recorded hourly using a Synergy 2 Multi-Mode reader (BioTek, Winooski, VT). The RFU results were subtracted with the RFU readings from a control strain, which was of the same genetic background but without the *gfp* promoter fusion, before normalization against the corresponding OD_600_ of the culture (89).

### Construction of genetic mutants

Genetic mutants of *S. mutans* were constructed through allelic exchange strategy and natural transformation facilitated by the use of competence-stimulating peptide (30). Recombinant DNA fragments used in this study were generated by PCR amplification and ligated together with antibiotic cassettes via Gibson Assembly (23). Primers used in DNA amplification and RT-qPCR are listed in Table S9. All mutants used in this study have been validated by PCR followed by Sanger sequencing. Promoter: reporter fusions were constructed using the plasmid pMC340B by inserting a promoter of interest in front of a promoterless *gfp* (green fluorescent protein) or *cat* (chloramphenicol acetyltransferase) gene, followed by integration at a distal site (between *mtlA* and *phnA*) of the chromosome unrelated to functions studied here (20).

### RNA deep sequencing, transcriptomic analysis, and reverse transcriptase quantitative PCR (RT-qPCR).

To compare bacterial gene regulation in response to fructose metabolism and exposure to methylglyoxal, *S. mutans* strain UA159 was cultivated to early exponential phase (OD_600_ = 0.4) in a chemically defined medium FMC (88) supplemented with 20 mM glucose. Bacteria were harvested by centrifugation (3,800× *g*, room temperature, 10 min), resuspended in fresh FMC containing 50 mM fructose, 50 mM glucose, or 5 mM methylglyoxal. An equal portion of the culture was kept in the original FMC-glucose medium as control. All samples were returned to incubation for 30 min before being harvested for RNA extraction.

RNA extraction and deep sequencing were carried out by following a previously published protocol detailed elsewhere (90). Briefly, cells were disrupted using a beadbeater in the presence of equal volume of acidic phenol:chloroform. After centrifugation, the cell lysate was processed using the RNeasy minikit (Qiagen, Germantown, MD) and an RNase-free DNase I solution (Qiagen) for extraction of total RNA. The RNA samples were shipped on dry ice to SeqCenter (Pittsburgh, PA) for quality check, cDNA preparation, and deep sequencing, yielding up to 12 million paired end reads (2 × 51 bp) per sample. The analysis of the RNA-seq data was performed in software R version 3.5.2 Eggshell Igloo as described previously (90). The compositional matrix of the expression data was normalized with the voom (91) function from the R package limma (92). The statistical analysis of the expression data (Table S10) was conducted using the package edgeR (version 3.24.3). A false-discovery rate (FDR) of 0.05 and a fold-of-change of 2.0 were used as the cutoff values for identification of genes with differential expression, consistent with the published work on H_2_O_2_ transcriptome in *S. mutans* (36). Findings from this analysis were validated by reverse transcriptase quantitative PCR (RT-qPCR) (30) carried out on 20 selected genes, following a previously published protocol detailed elsewhere (93, 94). The sequencing data from this study have been deposited at NCBI Gene Expression Omnibus (GEO) under the accession numbers GSE279080.

For comparative analysis under different treatments, we obtained the RNA-seq datasets (GEO, GSE98526) of an analysis previously performed on strain UA159 that was treated with 0.5 mM H_2_O_2_ for 5 min (36). Comparison between transcriptomes was carried out in R statistical language using the package edgeR as done previously (95).

### Long-term persistence assay

For comparison of the ability to persist under low-pH and starvation conditions, the cultures of *S. mutans* were each diluted 1000-fold into an FMC medium supplemented with 20 mM glucose, 20 mM fructose, or 19 mM glucose and 1 mM fructose combined. The cultures were incubated at 37°C in an ambient atmosphere supplemented with 5% CO_2_ for a duration of 11 days. After one or two days of incubation, a 30-second sonication at 100% power (FB120 water bath sonicator, Fisher Scientific) was applied to all cultures to disperse bacterial chains and aggregates, then a 100-µL aliquot of each culture was removed. The cultures were returned to incubation without any other treatments, and the 100-µL aliquots were each diluted decimally in sterile PBS and plated on BHI agar. After 2 days of incubation, the CFU on each plate were enumerated. Plating was repeated at specified time points thereafter till end of the experiment.

### Mixed-species competition in planktonic cultures

For interspecies competition in liquid cultures, differentially marked strains of *S. mutans* (UA159-Km and Δ*fruI::Km*) (96) and *S. sanguinis* (MMZ1945, Em resistant) (97) were each cultured in BHI to the exponential phase (OD_600_ = 0.5). An inoculum of approximately 10^6^ CFU/ml of *S. mutans* together with similar numbers of *S. sanguinis* were added to FMC supplemented with 20, or 200 mM glucose or fructose, then cultured for 20 h in a 5%-CO_2_ environment at 37°C. At both the start and the end of the incubation, cultures were sonicated for 15 sec, decimally diluted and plated on respective antibiotic plates to enumerate CFU of both species. The competitive index was calculated by following this formula: [SMU(*t*_end_)/SSA(*t*_end_)] / [SMU(*t*_start_)/SSA(*t*_start_)], with values >1 indicating SMU (*S. mutans*) being more competitive than SSA (*S. sanguinis*), and vice versa.

### pH drop and chloramphenicol acetyltransferase (CAT) assay

Glycolytic profiling via pH drop (98) and CAT assay (99) were conducted according to previously published protocols detailed elsewhere.

### Statistics

Statistical analysis of data was carried out using the software of Prism (GraphPad of Dotmatics, San Diego, CA)

## Acknowledgements

This study was supported by a grant DE12236 from NIDCR and a startup fund from University of Florida to LZ. JL was supported by grants DE032555 and DE019783 from NIDCR. Parts of Fig. 8 were created using BioRender.com.

