## Supplemental tables S1, S7, S8, S9 and Figures S1 to S6 for "FRUCTOSE ACTIVATES A STRESS RESPONSE SHARED BY METHYLGLYOXAL AND HYDROGEN PEROXIDE IN *STREPTOCOCCUS MUTANS*"

**Table S1.** Genes differentially expressed after treatments with methylglyoxal (MG), fructose (Fru), and glucose (Glc) relative to untreated controls. Results of $H_2O_2$ treatment was obtained from a previous study (1). These genes are discussed in the manuscript and organized based on functional categories. See Table S2 for the complete list. Red and green indicate increased and decreased expression, respectively, with shades denoting degrees of change.

| SMU Number | Gene Name | Gene Function | MG | Fru | Glc | $H_2O_2$ |
|---|---|---|---|---|---|---|
| ***Thiol homeostasis/oxidative stress*** | | | | | | |
| 140 | *gshR* | glutathione reductase | | 3.53 | 4.09 | |
| 141 | *SMU_141* | conserved hypothetical protein | 0.63 | 3.78 | 4.08 | |
| 267 | *gshAB* | glutamate--cysteine ligase (gamma ECS) | 1.14 | 1.29 | 1.00 | |
| 382 | *SMU_382c* | possible oxidoreductase | 3.67 | | | |
| 383 | *SMU_383c* | undetermined reductase or epimerase | 3.73 | | | |
| 463 | *trxB* | thioredoxin reductase (NADPH) | 1.90 | 1.31 | | 1.75 |
| 629 | *sodA* | superoxide dismutase | 3.00 | 2.89 | 1.02 | 2.22 |
| 667 | *nrdB* | ribonucleotide reductase, small subunit | 3.50 | 1.31 | 0.66 | 1.54 |
| 668 | *nrdA* | ribonucleotide reductase, large subunit | 3.07 | 1.09 | | 1.37 |
| 669 | *nrdH* | glutaredoxin | 2.63 | 0.83 | -0.63 | 1.48 |
| 676 | *gapN* | NADP-dependent G3P dehydrogenase | 1.21 | 1.20 | 0.93 | 1.40 |
| 677 | *rmeA* | transcriptional regulator, MerR family | 2.46 | 1.01 | | |
| 678 | *ycgG* | aldo/keto oxidoreductase | 3.15 | 1.21 | 0.96 | |
| 679 | *ycgG* | aldo/keto oxidoreductase | 3.24 | 1.49 | 1.12 | |
| 680 | *SMU_680* | carboxymuconolactone decarboxylase family protein | 3.20 | 1.22 | 0.71 | |
| 681 | *SMU_681* | conserved hypothetical protein | 3.14 | 1.14 | | |
| 728 | *SMU_728* | oxidoreductase | | 1.03 | | |
| 764 | *ahpC* | alkyl hydroperoxide reductase subunit C | 3.76 | 2.18 | | 2.33 |
| 765 | *ahpF* | alkyl hydroperoxide reductase subunit F | 4.22 | 2.30 | | 2.24 |
| 837 | *ycgG* | aldo/keto oxidoreductase | | | 1.39 | |
| 838 | *gor* | glutathione reductase | 3.44 | 1.51 | 1.17 | 2.14 |
| 924 | *tpx* | thiol peroxidase | 3.01 | 2.48 | 1.06 | 2.72 |
| 991 | *SMU_991* | possible ribonucleotide reductase | 1.62 | 1.26 | | 1.93 |
| 1040 | *ydfG* | oxidoreductase, short-chain dehydrogenase/reductase | 1.34 | | | |
| 1117 | *nox* | $H_2O$-forming NADH Oxidase | 1.81 | 1.42 | 0.97 | 1.94 |
| 1296 | *gst* | glutathione S-transferase | 2.70 | 2.43 | 1.46 | 1.79 |
| 1473 | *SMU_1473c* | oxidoreductase, short-chain dehydrogenase/reductase family | 1.21 | 1.05 | | |
| 1602 | *frp* | NAD(P)H-flavin oxidoreductase | 1.78 | 1.19 | 1.16 | |
| 1603 | *lguL* | lactoylglutathione lyase | 2.14 | 1.78 | 1.70 | |

| SMU Number | Gene Name | Gene Function | MG | Fru | Glc | $H_2O_2$ |
|---|---|---|---|---|---|---|
| 1869 | *trxA* | thioredoxin | 1.63 | 2.28 | 1.34 | 2.25 |
| 2070 | *SMU_2070* | conserved hypothetical protein | 3.21 | | | |
| 2071 | *nrdG* | ribonucleoside-triphosphate reductase activating protein | 3.66 | | | |
| 2072 | *SMU_2072c* | possible acetyltransferase, GNAT family | 3.70 | | | -1.22 |
| 2073 | *SMU_2073c* | conserved hypothetical protein | 3.59 | -1.09 | -1.61 | |
| 2074 | *nrdD* | anaerobic ribonucleoside-triphosphate reductase | 3.52 | -1.40 | | |
| ***Metal homeostasis*** | | | | | | |
| 182 | *sloA* | ABC transporter for iron/manganese | -1.87 | -2.89 | -1.51 | -1.01 |
| 183 | *sloB* | manganese ABC transporter permease element | -0.92 | -1.20 | | -0.96 |
| 184 | *sloC* | ABC transporter element, iron/manganese binding protein | | | -1.34 | -1.14 |
| 185 | *SMU_185* | hypothetical protein | | -0.60 | -1.51 | -1.56 |
| 186 | *sloR* | metal-dependent transcriptional regulator | -0.77 | | | -1.01 |
| 247 | *sufC* | ABC transporter, ATP-binding protein | 2.24 | 0.98 | 0.72 | 1.89 |
| 248 | *sufD* | ABC transporter permease | 2.56 | 1.22 | 1.09 | 1.82 |
| 249 | *sufS* | class-V aminotransferase, NifS protein homolog, | 2.83 | 1.31 | 0.96 | 1.80 |
| 250 | *sufU* | nitrogen fixation-like protein, NifU | 2.91 | 1.01 | | 1.90 |
| 251 | *sufB* | ABC transporter permease | 2.76 | 1.08 | 0.77 | 1.70 |
| 424 | *copY* | negative transcriptional regulator, CopY | 3.82 | 0.84 | 1.23 | 1.24 |
| 426 | *copA* | copper-transporting ATPase | 3.72 | 0.64 | 0.77 | 1.18 |
| 427 | *copZ* | copper chaperone | 3.47 | 0.82 | | 1.09 |
| 540 | *dpr* | peroxide resistance protein/iron binding protein | 2.96 | 1.87 | | 1.99 |
| 569 | *feoA* | ferrous ion transport protein A | -2.44 | | | -1.17 |
| 570 | *feoB* | ferrous ion transport protein B | -1.65 | -0.66 | 0.77 | -1.00 |
| 1561 | *trkB* | potassium uptake protein B | 2.20 | 1.64 | | |
| 1562 | *trkA* | potassium uptake protein A | 2.32 | 1.53 | | |
| 1993 | *adcB* | zinc ABC transporter, permease | | | | 1.08 |
| 1994 | *adcC* | zinc ABC transport, ATP-binding protein | | | | 1.10 |
| 1995 | *adcR* | zinc transport transcriptional repressor | | | | 1.04 |
| 2057 | *zccE* | zinc/cadmium-efflux ATPase | -0.42 | 2.44 | 1.17 | |
| ***DNA and protein repair*** | | | | | | |
| 80 | *hrcA* | heat-inducible transcription repressor | -2.47 | | | |
| 81 | *grpE* | co-chaperone protein GrpE | -2.12 | | | |
| 82 | *dnaK* | chaperone protein, DnaK | -1.33 | | | |
| 561 | *mutT* | mutator protein, Nudix hydrolase | 2.46 | | | |
| 562 | *clpE* | ATP-dependent protease | 2.86 | 1.44 | | 1.42 |
| 809 | *uvrB* | excinuclease ABC subunit B | | | 1.26 | |
| 956 | *clpL* | ATP-dependent Clp protease | 1.77 | 1.13 | | 1.11 |
| 1649 | *exoA* | exodeoxyribonuclease III | 1.68 | 1.19 | 1.10 | 1.93 |
| 1650 | *end3* | endonuclease III | 1.82 | | | 1.56 |
| 1850 | *pepP* | XAA-Pro aminopeptidase | 1.78 | 1.16 | 0.89 | 1.45 |

| SMU Number | Gene Name | Gene Function | MG | Fru | Glc | $H_2O_2$ |
|---|---|---|---|---|---|---|
| 1851 | *uvrA* | excinuclease ABC subunit A | 1.34 | 1.03 | | 1.70 |
| 1865 | *mutY* | A/G-specific adenine glycosylase | 2.41 | 1.24 | 0.66 | 1.66 |
| 2029 | *clpC* | ATP-dependent Clp protease | 1.11 | | | |
| 2036 | *pepO* | endopeptidase O | 1.61 | 1.18 | 1.07 | 1.42 |
| 2085 | *recA* | recombinase A | | 1.18 | | |
| ***Transcription regulators*** | | | | | | |
| 61 | *comR* | transcriptional regulator | 1.01 | | | 1.55 |
| 363 | *glnR* | glutamine synthetase repressor | | -2.29 | | |
| 381 | *plcR* | transcriptional regulator | 1.45 | | | |
| 486 | *liaS* | two-component sensor histidine kinase | | | | -1.00 |
| 487 | *liaR* | two-component response regulator | | | | -1.01 |
| 576 | *lytT* | response regulator | -1.55 | -1.58 | | |
| 577 | *lytS* | sensor histidine kinase | -2.26 | -1.51 | | |
| 593 | *perR* | ferric uptake regulator protein | 1.06 | 1.08 | | 1.03 |
| 660 | *spaK* | histidine kinase | | -1.05 | | |
| 661 | *spaR* | transcriptional regulator | 1.72 | | | |
| 928 | *relS* | sensor histidine kinase | 1.00 | | | |
| 1037 | *phoR* | histidine kinase | 2.12 | | | |
| 1038 | *ycbL* | response regulator | 1.79 | | | |
| 1065 | *nagR* | transcriptional regulator, GntR family | -1.42 | | | |
| 1097 | *SMU_1097c* | transcriptional regulator | 1.37 | | -1.42 | |
| 1226 | *SMU_1226c* | histidine protein kinase | 1.03 | | | |
| 1287 | *pmrA* | transcriptional regulator | -1.03 | | -1.48 | 1.12 |
| 1398 | *irvR* | phage associated repressor protein | -1.60 | | | |
| 1515 | *vicX* | *gtfB/C* regulator | 1.28 | | | |
| 1516 | *vicK* | two-component sensor histidine kinase | 1.17 | | | |
| 1814 | *scnK* | histidine kinase, ScnK homolog | 2.50 | 1.37 | | 1.58 |
| 1815 | *scnR* | response regulator, ScnR homolog | 2.26 | 1.37 | 0.59 | 1.72 |
| 1854 | *hdrR* | high density responsive regulator | | 1.35 | | |
| 1915 | *comC* | competence stimulating peptide | 2.15 | | | |
| 1916 | *comD* | histidine kinase | 2.95 | 1.01 | | 1.24 |
| 1917 | *comE* | response regulator | 2.78 | | | 1.31 |
| 1924 | *gcrR* | response regulator for glucan-binding protein | | -1.77 | -1.09 | |
| 1963 | *levQ* | sugar-binding periplasmic protein | | -1.02 | -1.23 | -1.07 |
| 1964 | *levR* | two-component response regulator | | | -1.07 | -1.20 |
| 1965 | *levS* | histidine kinase | | | | -1.12 |
| 1966 | *levT* | ABC transport ribose-binding protein, periplasmic | | | | -1.19 |
| 2027 | *lexA* | transcriptional regulator/repressor | 3.03 | 3.08 | 1.09 | 1.29 |
| 2084 | *spxA2* | oxidative stress regulator | | 1.30 | | |
| ***Osmoprotection*** | | | | | | |
| 1057 | *satE* | acid tolerance protein | | 1.05 | | |
| 1058 | *satD* | acid tolerance protein | | 1.26 | | |
| 1060 | *ffh* | signal recognition particle | | 1.13 | | |

| SMU Number | Gene Name | Gene Function | MG | Fru | Glc | $H_2O_2$ |
|---|---|---|---|---|---|---|
| 1061 | *ylxM* | DNA-binding protein | | 1.22 | | |
| 1062 | *busAB* | glycine-betaine binding ABC transporter permease | 1.09 | 1.57 | | |
| 1063 | *opuAA* | amino acid ABC transporter, ATP-binding protein | | 1.68 | | |
| 2116 | *opuCa* | ABC transporter, ATP-binding protein | | 1.05 | | |
| 2117 | *opuCb* | ABC transporter permease | | 1.12 | | |
| 2118 | *opuCc* | ABC transporter, substrate-binding protein | | 1.13 | | |
| 2119 | *opuCd* | ABC transport permease | | 1.07 | | |
| ***F1F0 ATPase*** | | | | | | |
| 1527 | *atpA* | ATPase epsilon subunit | | -1.06 | | |
| 1528 | *atpB* | ATPase beta subunit | | -1.13 | | |
| 1529 | *atpC* | ATPase gamma subunit | | -1.31 | | |
| 1530 | *atpD* | ATPase alpha subunit | | -1.20 | | |
| 1531 | *atpE* | ATPase delta subunit | | -1.61 | -1.25 | |
| 1532 | *atpF* | ATPase b subunit | | -1.35 | | |
| 1533 | *atpG* | ATPase a subunit | | -1.40 | | |
| 1534 | *atpH* | ATPase c subunit | | -1.26 | | |
| ***Fatty acid biosythesis*** | | | | | | |
| 1735 | *accD* | acetyl-CoA carboxylase beta subunit | 0.72 | -1.15 | | |
| 1736 | *accC* | acetyl-CoA carboxylase biotin carboxylase subunit | 0.61 | -1.29 | | |
| 1737 | *fabZ* | (3R)-hydroxymyristoyl-(acyl carrier protein) dehydratase | 0.77 | -1.38 | | |
| 1738 | *accB* | biotin carboxyl carrier protein of acetyl-CoA carboxylase | 0.80 | -1.46 | | |
| 1739 | *fabF* | 3-oxoacyl-(acyl-carrier-protein) synthase | 0.69 | -1.51 | | |
| 1740 | *fabG* | 3-oxoacyl-acyl-carrier-protein reductase | | -1.78 | | |
| 1741 | *fabD* | [acyl-carrier-protein] S-malonyltransferase | | -1.74 | | |
| 1742 | *fabK* | trans-2-enoyl-ACP reductase II | | -1.56 | | |
| 1743 | *acp* | acyl carrier protein | | -1.55 | -1.56 | |
| 1744 | *fabH* | 3-oxoacyl-[acyl-carrier-protein] synthase III | -0.76 | -1.39 | | |
| 1745 | *fabT* | transcriptional regulator, MarR family | -1.11 | -1.06 | -0.66 | |
| ***Other stress/competence-related proteins*** | | | | | | |
| 20 | *mreC* | cell shape-determining protein MreC | -2.19 | -1.36 | -1.45 | |
| 21 | *mreD* | cell shape-determining protein MreD | -1.95 | -2.02 | -0.59 | |
| 150 | *nlmA* | non-lantibiotic mutacin IV A | | | | -1.30 |
| 151 | *nlmB* | non-lantibiotic mutacin IV B | | | | -1.13 |
| 152 | *SMU_152* | immunity protein for NlmAB | | | -1.88 | -1.64 |
| 153 | *SMU_153* | hypothetical protein | | | | -1.75 |
| 284 | *nlmT* | transport and processing of non-lantibiotic mutacins | | -1.24 | -1.38 | -1.05 |

| SMU Number | Gene Name | Gene Function | MG | Fru | Glc | $H_2O_2$ |
|---|---|---|---|---|---|---|
| 285 | *nlmE* | transport and processing of non-lantibiotic mutacins | | -1.13 | -1.16 | -1.16 |
| 474 | *luxS* | S-ribosylhomocysteine lyase (S-ribosylhomocysteinase) | -1.34 | | | |
| 503 | *SMU_503c* | hypothetical protein | -1.86 | -1.69 | -1.58 | -1.11 |
| 921 | *rcrR* | transcriptional regulator | 1.35 | | -1.31 | 1.21 |
| 922 | *rcrP* | ABC transporter, ATPase component | 1.75 | | | 1.07 |
| 923 | *rcrQ* | AB transporter, ATPase component | 1.87 | | | 1.08 |
| 1046 | *relQ* | GTP pyrophosphokinase | -1.08 | 1.03 | | |
| 574 | *lrgB* | effector of murein hydrolase | -2.42 | -2.46 | -1.11 | |
| 575 | *lrgA* | murein hydrolase regulator | -2.68 | -3.00 | -1.63 | |
| 1700 | *cidB* | LrgB-like protein; possible murein hydrolase regulator | 1.93 | 1.39 | | 1.31 |
| 1701 | *cidA* | conserved hypothetical protein | 1.90 | 1.05 | | 1.29 |
| 1702 | *pgpB* | uncharacterized phosphatase | 1.73 | 1.82 | | 1.59 |
| 1703 | *SMU_1703c* | conserved hypothetical protein | 1.41 | 1.78 | | 1.75 |
| 1980 | *SMU_1980c* | conserved hypothetical protein | 3.14 | 2.29 | 1.08 | |
| 1981 | *comGF* | competence protein G | 3.42 | 2.29 | | |
| 1982 | *SMU_1982c* | conserved hypothetical protein | 2.76 | 1.96 | | |
| 1983 | *comYD* | late competence protein | 2.20 | | | |
| 1984 | *comYC* | late competence protein | 2.50 | 1.26 | | |
| 1985 | *comYB* | late competence protein | 2.19 | | | |
| 1987 | *comYA* | late competence protein | 2.28 | | | |
| 1988 | *SMU_1988c* | probable DNA binding protein | 2.80 | 1.43 | | 2.19 |
| 1997 | *comX* | competence-specific sigma factor | -1.57 | | -1.46 | |
| 2044 | *relA* | GTP pyrophosphokinase | | | | 1.22 |
| ***Carbohydrate metabolism*** | | | | | | |
| 99 | *fbaA* | fructose-bisphosphate aldolase | | -1.25 | | |
| 100 | *nigB* | nigerose PTS system, IIB component | | -1.52 | | |
| 105 | *nigR* | nigerose operon transcriptional repressor | | -1.15 | | |
| 113 | *pfk* | fructose-1-phosphate kinase | | 1.39 | 1.13 | |
| 114 | *fruC* | PTS system, fructose-specific IIBC component | | | 1.08 | |
| 119 | *adh* | alcohol dehydrogenase class III | 1.02 | 1.74 | 1.76 | |
| 127 | *adhA* | acetoin dehydrogenase E1 component alpha subunit | 2.76 | | | 1.72 |
| 128 | *adhB* | acetoin dehydrogenase E1 component beta subunit | 3.33 | | | 1.88 |
| 129 | *adhC* | dihydrolipoamide S-acetyltransferase | 3.50 | | | 1.91 |
| 130 | *adhD* | dihydrolipoamide dehydrogenase | 3.30 | 1.27 | | 1.96 |
| 131 | *lplA* | lipoate-protein ligase | 3.43 | 1.39 | | 1.88 |
| 135 | *mleR* | transcriptional regulator | -1.05 | | | |
| 137 | *mleS* | malolactic enzyme | 0.62 | 4.00 | 4.36 | |
| 138 | *mleP* | malate permease/auxin efflux carrier | | 3.78 | 4.08 | |
| 139 | *oxdC* | oxalate decarboxylase | | 3.80 | 4.33 | |
| 148 | *adhE* | alcohol-acetaldehyde dehydrogenase | -1.11 | | 1.26 | |

| SMU Number | Gene Name | Gene Function | MG | Fru | Glc | $H_2O_2$ |
|---|---|---|---|---|---|---|
| 270 | *sgaT* | ribulose monophosphate PTS pathway enzyme IIC | | | 1.36 | |
| 271 | *ptxB* | L-ascorbate-specific PTS enzyme IIB component | 1.88 | 1.00 | 0.96 | 1.19 |
| 272 | *ptxA* | L-ascorbate-specific PTS enzyme IIA component | 2.13 | 1.04 | 1.03 | 0.90 |
| 273 | *rmpD* | hexulose-6-phosphate synthase | 2.30 | 1.11 | 0.87 | 1.01 |
| 274 | *rmpE* | hexulose-6-phosphate isomerase | 2.48 | 1.17 | 1.01 | 1.16 |
| 275 | *rmpF* | L-ribulose 5-phosphate 4-epimerase | 2.59 | 1.66 | 1.06 | 1.11 |
| 276 | *SMU_276c* | hypothetical protein | 2.37 | | -2.62 | |
| 311 | *srlA* | sorbitol PTS system enzyme IIC2 | | 1.02 | | |
| 312 | *srlE* | sorbitol PTS system enzyme IIBC | | 1.36 | | |
| 313 | *srlB* | sorbitol PTS system enzyme IIA | | 1.68 | | |
| 402 | *pfl* | pyruvate formate-lyase | | -1.49 | | |
| 507 | *sppR* | transcriptional regulator, DeoR family | | 4.56 | | |
| 508 | *sppA* | hexose-phosphate phosphohydrolase | | 4.56 | | |
| 674 | *ptsH* | PTS EI | -2.25 | | | |
| 675 | *ptsI* | PTS HPr | -1.14 | | | |
| 754 | *hprK* | HPr (serine) kinase/phosphatase | | 1.34 | 1.15 | |
| 870 | *fruR* | transcriptional repressor of the fructose operon | | 1.68 | | |
| 871 | *fruK* | fructose-1-phosphate kinase | | 1.99 | | |
| 872 | *fruI* | fructose-specific PTS system enzyme IIBC component | | 1.87 | | |
| 876 | *msmR* | MSM operon regulatory protein | -1.93 | -1.26 | | -1.28 |
| 877 | *agaL* | alpha-galactosidase | -1.46 | -1.73 | | -1.36 |
| 878 | *msmE* | ABC transporter, sugar-binding protein | | -1.79 | | -1.25 |
| 879 | *msmF* | ABC transporter, sugar permease protein | | -1.95 | | |
| 880 | *msmG* | ABC transporter permease | | -1.78 | | |
| 881 | *gtfA* | sucrose phosphorylase | | -1.64 | | |
| 882 | *msmK* | ABC transporter ATP-binding protein | | -1.54 | | |
| 883 | *dexB* | glucan 1,6-alpha-glucosidase | | -1.40 | | |
| 886 | *galK* | galactokinase | | -1.46 | | |
| 910 | *gtfD* | glucosyltransferase-S | | 1.24 | 1.82 | |
| 1004 | *gtfB* | glucosyltransferase-I | | | 1.82 | |
| 1005 | *gtfC* | glucosyltransferase-SI | | -1.10 | | |
| 1184 | *mtlR* | transcriptional regulator | -1.05 | | | |
| 1185 | *mtlA* | mannitol PTS EII | -1.73 | | | |
| 1396 | *gbpC* | glucan-binding protein C | 2.39 | 1.32 | 1.27 | 1.28 |
| 1451 | *aldB* | alpha-acetolactate decarboxylase | 2.86 | | | 1.66 |
| 1452 | *alsS* | alpha-acetolactate synthase | 2.33 | | | 1.54 |
| 1489 | *lacX* | aldose 1-epimerase | | 1.13 | | |
| 1492 | *lacF* | PTS system, lactose-specific IIA component | | 1.85 | 1.92 | |
| 1493 | *lacD* | tagatose-1,6-bisphosphate aldolase | | 1.39 | | |
| 1494 | *lacC* | tagatose-6-phosphate kinase | | 1.59 | 0.65 | |

| SMU Number | Gene Name | Gene Function | MG | Fru | Glc | $H_2O_2$ |
|---|---|---|---|---|---|---|
| 1495 | *lacB* | galactose-6-phosphate isomerase | -0.78 | 1.59 | 0.98 | |
| 1496 | *lacA* | galactose-6-phosphate isomerase | | 1.71 | 1.18 | |
| 1498 | *lacR* | lactose phosphotransferase system repressor | | 3.11 | | |
| 1537 | *glgD* | glycogen biosynthesis protein | | -1.06 | | |
| 1564 | *glg* | glycogen phosphorylase | | -1.24 | | |
| 1565 | *malM* | 4-alpha-glucanotransferase | | -1.02 | | |
| 1566 | *malR* | maltose operon transcriptional repressor | 1.53 | | | 1.57 |
| 1568 | *malE* | maltose / maltodextrin-binding protein | -1.55 | -2.32 | | |
| 1569 | *malF* | maltodextrin ABC transport system permease | | -2.00 | | |
| 1570 | *malG* | maltose / maltodextrin ABC transport system (permease) | | -2.06 | | |
| 1571 | *msmK* | ABC-type transport system ATP-binding protein (maltose) | | -1.89 | | |
| 1596 | *celD* | cellobiose PTS system IIC | -1.59 | -2.92 | -1.02 | |
| 1597 | *celX* | conserved hypothetical protein | -2.21 | -3.06 | -2.52 | |
| 1598 | *celC* | cellobiose PTS system IIA component | -2.43 | -3.00 | -1.11 | |
| 1599 | *celR* | cellobiose transcriptional regulator | -2.80 | -3.62 | -1.71 | |
| 1600 | *celB* | cellobiose PTS system IIB | -3.15 | -2.59 | -1.18 | |
| 1664 | *acoB* | acetoin utilization protein, acetoin dehydrogenase | 1.66 | 0.76 | | |
| 1692 | *pflA* | pyruvate-formate lyase activating enzyme | 1.36 | | | 1.81 |
| 1841 | *scrA* | sucrose PTS system EIIABC components | -1.99 | -2.13 | | |
| 1843 | *scrB* | sucrose-6-phosphate hydrolase | -1.65 | | | |
| 1844 | *scrR* | sucrose operon repressor | -1.07 | -1.24 | | |
| 1867 | *adhB* | alcohol dehydrogenase | 1.75 | 2.12 | 1.97 | 1.85 |
| 1877 | *manL* | mannose PTS system component IIAB | -2.50 | -2.79 | -0.85 | |
| 1878 | *manM* | mannose PTS system component IIC | -1.76 | -2.34 | | |
| 1879 | *manN* | mannose PTS system component IID | -1.24 | -2.25 | -0.92 | |
| 1956 | *levX* | conserved hypothetical protein | | 2.07 | | |
| 1957 | *levG* | fructose-specific Enzyme IID component | | 2.16 | | |
| 1958 | *levF* | fructose-specific Enzyme IIC component | | 2.15 | | |
| 1960 | *levE* | fructose-specific Enzyme IIB component | | 1.58 | | |
| 1961 | *levD* | fructose-specific Enzyme IIA component | | 1.77 | | |
| 2028 | *ftf* | fructosyltransferase | | 1.48 | 1.72 | |
| 2047 | *malT* | maltose PTS enzyme II | | -1.71 | | |
| ***Amino acid metabolism*** | | | | | | |
| 261 | *aguR* | transcriptional regulator | 1.42 | 1.65 | | |
| 262 | *argF* | ornithine carbamoyltransferase | 1.48 | 2.21 | 1.16 | |
| 263 | *aguD* | amino acid permease /putrescine antiporter | 1.49 | 2.13 | 1.41 | |

| SMU Number | Gene Name | Gene Function | MG | Fru | Glc | $H_2O_2$ |
|---|---|---|---|---|---|---|
| 264 | *aguA* | agmatine deiminase | | 1.74 | 1.48 | |
| 265 | *aguC* | carbamate kinase | | 1.18 | | |
| 563 | *argF* | ornithine carbamoyltransferase | | 1.00 | | |
| 663 | *argC* | N-acetyl-gamma-glutamyl-phosphate reductase | 3.25 | 2.41 | 1.17 | |
| 664 | *argJ* | ornithine acetyltransferase / N-acetylglutamate synthase | 3.25 | 2.42 | 1.94 | |
| 665 | *argB* | acetylglutamate kinase | 3.32 | 2.24 | 1.68 | |
| 666 | *argD* | N-acetylornithine aminotransferase | 2.94 | 1.54 | 1.78 | 1.09 |
| 1665 | *livF* | branched chain amino acid ABC transporter, ATP-binding protein | 1.57 | 0.73 | | 1.05 |
| 1666 | *livG* | branched chain amino acid ABC transporter, ATP-binding protein | 1.54 | 0.71 | 0.14 | 1.16 |
| 1667 | *livM* | branched chain amino acid ABC transporter, permease | 1.35 | | | 1.29 |
| 1668 | *livH* | branched chain amino acid ABC transporter, permease | 1.36 | | | 1.24 |
| 1669 | *livK* | branched-chain amino acid ABC transporter, substrate-binding protein | 1.06 | | | 1.34 |
| 2097 | *ahrC* | arginine transcriptional repressor | -1.22 | | | |
| ***Translation*** | | | | | | |
| 169 | *rplM* | 50S ribosomal protein L13 | -1.22 | -1.18 | | |
| 170 | *rpsI* | 30S ribosomal protein S9 | | -1.32 | | |
| 697 | *infC* | translation initiation factor IF-3 | -1.06 | | | |
| 818 | *rpsU* | 30S ribosomal protein S21 | | -1.38 | | |
| 957 | *rplJ* | 50S ribosomal protein L10 | -1.80 | -1.89 | | |
| 960 | *rpl* | 50S ribosomal protein L7/L12 | -1.33 | -1.77 | | |
| 1127 | *rpsT* | 30S ribosomal protein S20 | -1.03 | | | |
| 1200 | *rpsA* | 30S ribosomal protein S1 | -1.21 | | | |
| 1288 | *rplS* | 50S ribosomal protein L19 | -1.06 | | | |
| 1627 | *rplK* | 50S ribosomal protein L11 | -1.11 | | | |
| 1860 | *rpsF* | 30S ribosomal protein S6 | -1.16 | | | |
| 2000 | *rplQ* | 50S ribosomal protein L17 | -1.66 | | | |
| 2002 | *rpsK* | 30S ribosomal protein S11 | -1.27 | | | |
| 2003 | *rpsM* | 30S ribosomal protein S13, N-terminal fragment | -1.06 | | | |
| 2003a | *rl36* | 50S ribosomal protein L36 | -1.05 | | -1.22 | |
| 2024 | *rplD* | 50S ribosomal protein L4, N-terminal fragment | | -1.01 | | |
| 2025 | *rplC* | 50S ribosomal protein L3 | | -1.06 | | |
| 2104a | *rl32* | 50S ribosomal protein L32 | -1.03 | | | |
| 2135 | *rpsD* | 30S ribosomal protein S4 | | -1.08 | -1.03 | |
| ***TnSmu1*** | | | | | | |
| 196 | *SMU_196c* | immunogenic secreted protein (transfer protein) | | 1.35 | 1.60 | |
| 197 | *SMU_197c* | hypothetical protein | | 1.09 | 1.35 | |
| 199 | *SMU_199c* | hypothetical protein | | 1.20 | 1.14 | |
| 200 | *SMU_200c* | hypothetical protein | | 1.98 | | |

| SMU Number | Gene Name | Gene Function | MG | Fru | Glc | $H_2O_2$ |
|---|---|---|---|---|---|---|
| 201 | *SMU_201c* | conserved hypothetical protein | | 1.12 | | |
| 202 | *SMU_202c* | conserved hypothetical protein | | 1.80 | 1.55 | |
| 204 | *SMU_204c* | hypothetical protein | | 1.80 | 1.57 | |
| 205 | *SMU_205c* | conserved hypothetical protein | | 1.91 | 1.28 | |
| 206 | *SMU_206c* | hypothetical protein | | 1.48 | 1.27 | |
| 207 | *SMU_207c* | transcriptional regulator | | | 1.53 | |
| 208 | *SMU_208c* | conserved hypothetical protein, FtsK/SpoIIIE family | | 1.01 | 1.46 | |
| 209 | *SMU_209c* | hypothetical protein | | 1.31 | 1.56 | |
| 210 | *SMU_210c* | hypothetical protein | | 1.15 | 1.33 | |
| 211 | *SMU_211c* | hypothetical protein | | 1.31 | 2.27 | |
| 212 | *SMU_212c* | hypothetical protein | | 1.20 | 1.50 | |
| 213 | *SMU_213c* | hypothetical protein | | 1.52 | 1.90 | |
| 214 | *SMU_214c* | hypothetical protein | | 1.27 | | |
| 215 | *SMU_215c* | hypothetical protein | | 1.72 | 1.09 | |
| 216 | *SMU_216c* | hypothetical protein | | 1.37 | | |
| 217 | *SMU_217c* | conserved hypothetical protein | | 1.34 | | |
| 218 | *SMU_218* | transcriptional regulator | | | -1.27 | |
| ***CRISPR-Cas*** | | | | | | |
| 1402 | *csn2* | conserved hypothetical protein | -1.28 | -2.56 | -1.95 | |
| 1403 | *cas2* | conserved hypothetical protein | -1.49 | -2.32 | -0.85 | |
| 1404 | *cas1* | conserved hypothetical protein | -1.30 | -2.24 | -1.22 | |
| 1405 | *cas9* | conserved hypothetical protein | -1.50 | -2.31 | -1.74 | |
| 1406 | *SMU_1406c* | possible oxidoreductase | -1.52 | -1.15 | | |
| 1753 | *cas2* | conserved hypothetical protein | | 1.59 | | 1.14 |
| 1754 | *cas1* | conserved hypothetical protein | | 1.91 | 1.09 | 1.29 |
| 1755 | *cas1* | conserved hypothetical protein | | 1.70 | | 1.21 |
| 1757 | *cas1* | conserved hypothetical protein | | 1.77 | | 1.04 |
| 1758 | *cas4* | conserved hypothetical protein | | 1.78 | | 1.36 |
| 1760 | *csd2* | conserved hypothetical protein | 1.03 | 2.02 | | 1.51 |
| 1761 | *cas8c* | conserved hypothetical protein | 1.48 | 1.79 | | 1.63 |
| 1762 | *csd1* | conserved hypothetical protein | 1.43 | 1.88 | | 1.87 |
| 1763 | *cas5* | conserved hypothetical protein | 1.47 | 1.94 | | 1.88 |
| 1764 | *cas3* | conserved hypothetical protein | 1.22 | 1.90 | | 2.09 |

**Table S7**. Expression of *fruRBA* in *S. sanguinis* SK36 (by RT-qPCR) and activities of the *fruR* promoter in UA159 (by CAT assay).

| | SK36 relative mRNA levels | | | UA159 P*fruR::cat* sp. act. |
|---|---|---|---|---|
| | *fruR* | *fruB* | *fruA* | |
| Control | 1.00 ± 0.07 | 1.00 ± 0.10 | 1.00 ± 0.06 | 186.18 ± 13.87 |
| 20 μM fructose | 9.70 ± 1.16 | 9.54 ± 3.15 | 7.98 ± 4.93 | 230.71 ± 7.22 |
| 50 mM fructose | 10.67 ± 1.74 | 10.29 ± 1.92 | 8.83 ± 3.22 | 271.29 ± 3.59 |
| 25% human sera | 1.27 ± 0.19 | 1.71 ± 0.54 | 2.04 ± 0.31 | ND |

Bacteria were cultured in FMC-glucose till exponential phase ($OD_{600}$ = 0.4) and then treated for 30 min with fructose or 25% pooled human sera. Human sera were used at 25% instead of 100% to maintain comparability with the other samples of the experiment. Gene expression was measured by RT-qPCR or CAT assays.

**Table S8.** pH and final $OD_{600}$ of 20-h TV cultures of *S. mutans* WT isolates (n = 4).

| Strains | Sugars | Resting pH | | | Final $OD_{600}$ | | |
|---|---|---|---|---|---|---|---|
| | | 20 mM | 100 mM | 200 mM | 20 mM | 100 mM | 200 mM |
| SMU_UA159 | Glucose | 4.92 | 4.93 | 4.93 | 0.83 | 0.72 | 0.57 |
| | Fructose | 4.76**** | 4.82** | 4.81*** | 0.82 | 0.73 | 0.64**** |
| SMU_UA101 | Glucose | 4.61 | 4.52 | 4.53 | 0.96 | 0.82 | 0.66 |
| | Fructose | 4.75*** | 4.55 | 4.61 | 0.86**** | 0.83 | 0.69* |
| SMU_GS-5 | Glucose | 4.73 | 4.65 | 4.64 | 0.93 | 0.80 | 0.64 |
| | Fructose | 4.62*** | 4.57** | 4.62 | 0.86** | 0.78 | 0.65 |
| SMU_10449 | Glucose | 4.81 | 4.73 | 4.73 | 0.95 | 0.85 | 0.70 |
| | Fructose | 4.82 | 4.70 | 4.77 | 0.77**** | 0.75**** | 0.64** |
| SMU_OMZ175 | Glucose | 4.62 | 4.53 | 4.53 | 1.00 | 0.84 | 0.69 |
| | Fructose | 4.75*** | 4.76**** | 4.86**** | 0.88**** | 0.74**** | 0.64*** |
| SMU_ST1 | Glucose | 4.77 | 4.72 | 4.75 | 0.86 | 0.74 | 0.61 |
| | Fructose | 4.88*** | 4.68 | 4.70 | 0.80*** | 0.76 | 0.69**** |
| SMU_SM6 | Glucose | 4.73 | 4.72 | 4.72 | 0.68 | 0.57 | 0.49 |
| | Fructose | 4.90*** | 4.88** | 5.03**** | 0.71 | 0.60 | 0.51 |

Results show average (standard deviation) of four independent cultures. Statistical significance was determined relative to glucose cultures of the same concentration using Two-Way ANOVA (*, $P$ <0.05; **, $P$ <0.01; ***, $P$ <0.001; ****, $P$ <0.0001). Color red denotes decreased value, and green for increased value, relative to glucose conditions.

**Table S9.** Primers used in this study.

| Primer name | Sequence | Purpose |
|---|---|---|
| fruI-1 | GGC GGT TTT ACT GGT AGG TT | Δ*fruI* |
| fruI-2GA | GCC ATT TAT TAT TTC CTT CCT CTT TTA GCC TGC AAG TCA AGC AAC A | Δ*fruI* |
| fruI-3GA | ATA TTT TAC TGG ATG AAT TGT TTT AGT AGA GCT GTT GGA GCA GTG ATT GCT | Δ*fruI* |
| fruI-4 | AGC TGC TTG GAG ATA ATC TTT ACC T | Δ*fruI* |
| SodpromF | CCGGAATTCGATGTAGCTTTGC | *sodA::gfp* |
| SodpromR | CGCGGATCCTTCCTCTTTTC | *sodA::gfp* |
| SMU.phoR-S | AAACACGGCAAAAATCAAGC | RT-qPCR |
| SMU.phoR-AS | TTGTCAGCTGATCCAAATGC | RT-qPCR |
| nlmA-S | TGGACAGCCAAACACTTTCA | RT-qPCR |
| nlmA-AS | CTGCTGAAGCAGTTGCACAT | RT-qPCR |
| Sm_sloA-AS | ACG AGC CAA GAG CAT ACG TT | RT-qPCR |
| Sm_sloA-S | TGG CAA TGT GTT CAA GAA GC | RT-qPCR |
| SMU_sloC-S | AAG GAA AAG TCG GGT GTC TG | RT-qPCR |
| SMU_sloC-AS | TTT AGC GGC AAT TGG GAT AC | RT-qPCR |
| celB-S | CAAAAACGCATTGAAGCAGA | RT-qPCR |
| celB-AS | GCTGCATCAGCCAACTGTTT | RT-qPCR |
| mreC-AS | TTT TGC CCC TCA AAT TTT TC | RT-qPCR |
| mreC-S | AGC GCT GAT GAG ATC GTA CA | RT-qPCR |
| SMU.508-S | ATG GGG CTC TGA TTG TTG AG | RT-qPCR |
| SMU.508-AS | GCG GAA ACT GCT TCT TGA TT | RT-qPCR |
| SMU.127-AS | GCA TAG AAC CAC CAC GAC CT | RT-qPCR |
| SMU.127-S | TTT TCA AAT CAC CGT GGA CA | RT-qPCR |
| ahpF-S | GAA ACC ATG GGT GGT CAA GT | RT-qPCR |
| ahpF-AS | GAG CCA TTA ATT GGG GTC CT | RT-qPCR |
| copY-S | ACC AGT CAG CGT CAA GGA AG | RT-qPCR |
| copY-AS | ACA AAT GCG CGA GAA AAC TT | RT-qPCR |
| SMU.667-S | TGA CTT GGA TGA CTG GCG TA | RT-qPCR |
| SMU.667-AS | TGG GAT TGC ATG GTG TCT AA | RT-qPCR |
| feoA-S | ACG GTT GAA ACA TTG CCT TT | RT-qPCR |
| feoA-AS | CGG CGC TTA GCT TCA TTA AC | RT-qPCR |
| SMUsloC-S | AAG GAA AAG TCG GGT GTC TG | RT-qPCR |
| SMU_sloC-AS | TTT AGC GGC AAT TGG GAT AC | RT-qPCR |
| gbpC-S | AGC TCA AAA GGC AGC TTA CG | RT-qPCR |
| gbpC-AS | TCCTTGGGCTTTTTCAACAC | RT-qPCR |
| scnK-S | CAAACGGCTGATTTCACAAG | RT-qPCR |
| scnK-AS | AAAGTGCGTTCCAATCTGCT | RT-qPCR |

| | | |
|---|---|---|
| SMU.924-S | TCAGTTGATTTGCCATTTGC | RT-qPCR |
| SMU.924-AS | CCATAAGCCTTTCCAAACGA | RT-qPCR |
| SMU_phoR-S | AAACACGGCAAAAATCAAGC | RT-qPCR |
| SMU_phoR-AS | TTGTCAGCTGATCCAAATGC | RT-qPCR |
| SMU_vicK-S | TTTGGCACAGGAGAAAAACC | RT-qPCR |
| SMU_vicK-AS | TAATCTTTCCGCTGCGATCT | RT-qPCR |
| SMU.679-S | TTG GTG CTT ATC GTG CTC TG | RT-qPCR |
| SMU.679-AS | TGG AAT AAC CGA CAC CGT TT | RT-qPCR |
| SMU.1602-S | GTC GCG GTT TAA AGG ACA AA | RT-qPCR |
| SMU.1602-AS | TAG CGC CAA ATC GTT TTC TT | RT-qPCR |
| gshAB-S | GAGGCCCTCCGCTATTATTC | RT-qPCR |
| gshAB-AS | TTGTGACTGGCTGCTGTTTC | RT-qPCR |
| SMU.1603-S | GAA GAC GAC CCC GAC TAT GA | RT-qPCR |
| SMU.1603-AS | TCG CTT CAA GGT CAT CAA CA | RT-qPCR |

**Fig. S1** Principal component analysis performed on transcriptomic datasets obtained from treatments of UA159 by 50 mM fructose (Fru), 5 mM methylglyoxal (MG), and the untreated control (Glc).

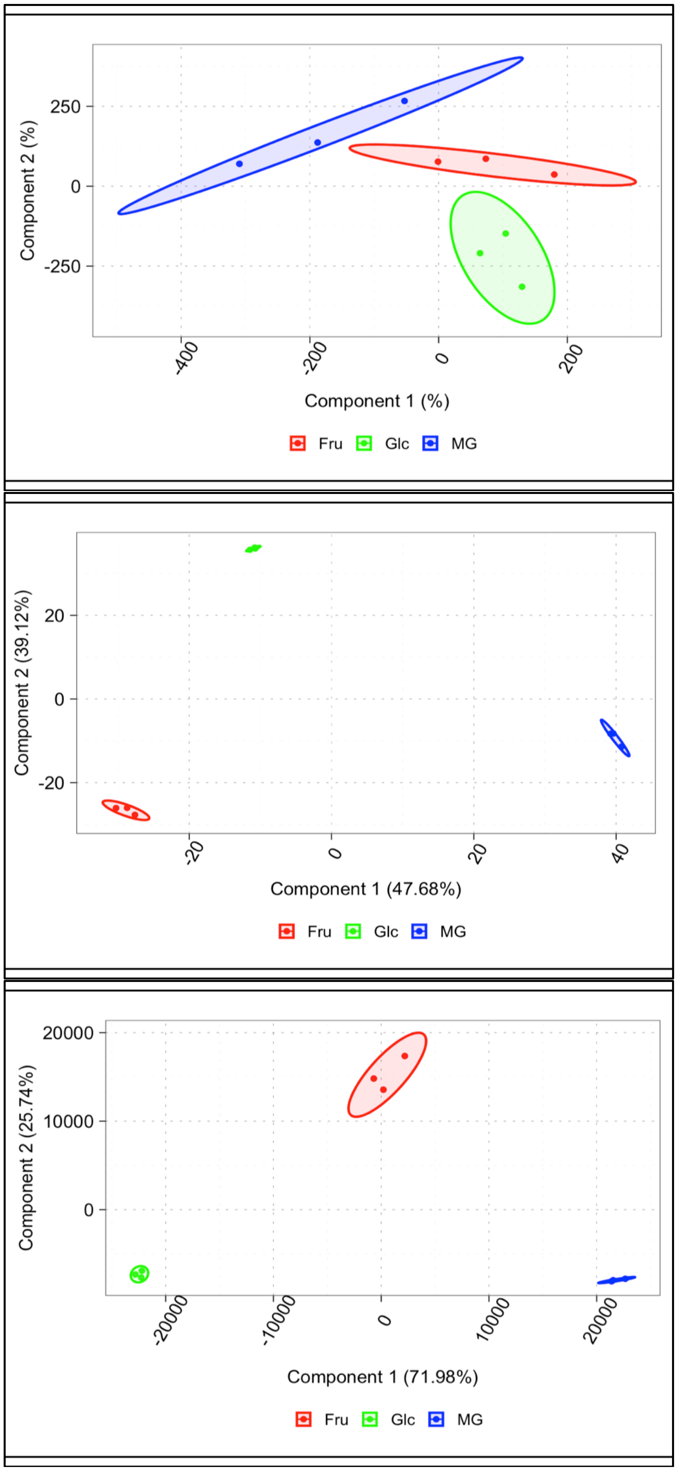

**Fig. S2** Gene Ontology (GO) term analyses performed on transcriptomes of UA159 impacted by (A) 50 mM fructose and (B) 5 mM methylglyoxal. Results shown are derived from molecular function analysis only. Blocks filled with blue indicate functions being repressed, whereas red colors indicate functions being activated by the treatment.

A

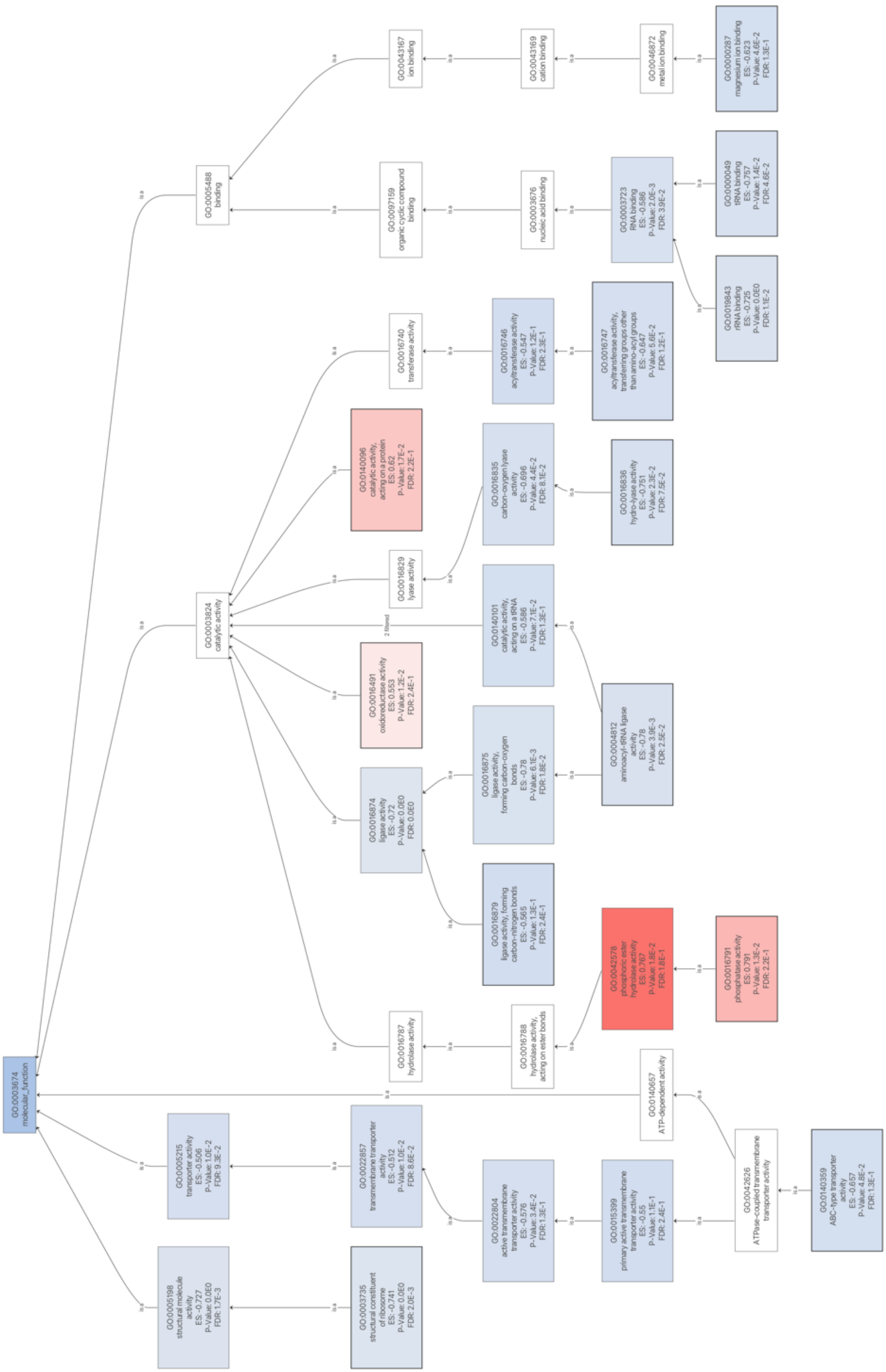
GO:0003674 molecular_function
GO:0005488 binding
GO:0003824 catalytic activity
GO:0005215 transporter activity ES:-0.506 P-Value: 1.0E-2 FDR: 9.3E-2
GO:0005198 structural molecule activity ES:-0.727 P-Value: 0.0E0 FDR: 1.7E-3
GO:0043167 ion binding
GO:0097159 organic cyclic compound binding
GO:0016740 transferase activity
GO:0140096 catalytic activity, acting on a protein ES: 0.62 P-Value: 1.7E-2 FDR: 2.2E-1
GO:0016829 lyase activity
GO:0016491 oxidoreductase activity ES: 0.553 P-Value: 1.2E-2 FDR: 2.4E-1
GO:0016874 ligase activity ES:-0.72 P-Value: 0.0E0 FDR: 0.0E0
GO:0016787 hydrolase activity
GO:0022857 transmembrane transporter activity ES:-0.512 P-Value: 1.0E-2 FDR: 8.6E-2
GO:0003735 structural constituent of ribosome ES:-0.741 P-Value: 0.0E0 FDR: 2.0E-3
GO:0043169 cation binding
GO:0003676 nucleic acid binding
GO:0016746 acyltransferase activity ES:-0.547 P-Value: 1.2E-1 FDR: 2.3E-1
GO:0016835 carbon-oxygen lyase activity ES:-0.696 P-Value: 4.4E-2 FDR: 8.1E-2
GO:0140101 catalytic activity, acting on a tRNA ES:-0.586 P-Value: 7.1E-2 FDR: 1.3E-1
GO:0016875 ligase activity, forming carbon-oxygen bonds ES:-0.78 P-Value: 6.1E-3 FDR: 1.8E-2
GO:0016879 ligase activity, forming carbon-nitrogen bonds ES:-0.565 P-Value: 1.3E-1 FDR: 2.4E-1
GO:0016788 hydrolase activity, acting on ester bonds
GO:0022804 active transmembrane transporter activity ES:-0.576 P-Value: 3.4E-2 FDR: 1.3E-1
GO:0046872 metal ion binding
GO:0003723 RNA binding ES:-0.586 P-Value: 2.0E-3 FDR: 3.9E-2
GO:0016747 acyltransferase activity, transferring groups other than amino-acyl groups ES:-0.647 P-Value: 5.6E-2 FDR: 1.2E-1
GO:0016836 hydro-lyase activity ES:-0.751 P-Value: 2.3E-2 FDR: 7.5E-2
GO:0004812 aminoacyl-tRNA ligase activity ES:-0.78 P-Value: 3.9E-3 FDR: 2.5E-2
GO:0042578 phosphoric ester hydrolase activity ES: 0.767 P-Value: 1.8E-2 FDR: 1.8E-1
GO:0140657 ATP-dependent activity
GO:0015399 primary active transmembrane transporter activity ES:-0.55 P-Value: 1.1E-1 FDR: 2.4E-1
GO:0000287 magnesium ion binding ES:-0.623 P-Value: 4.6E-2 FDR: 1.3E-1
GO:0000049 tRNA binding ES:-0.757 P-Value: 1.4E-2 FDR: 4.6E-2
GO:0019843 rRNA binding ES:-0.725 P-Value: 0.0E0 FDR: 1.1E-2
GO:0016791 phosphatase activity ES: 0.791 P-Value: 1.3E-2 FDR: 2.2E-1
GO:0042626 ATPase-coupled transmembrane transporter activity
GO:0140359 ABC-type transporter activity ES:-0.657 P-Value: 4.8E-2 FDR: 1.3E-1
is a
2 filtered

37

B

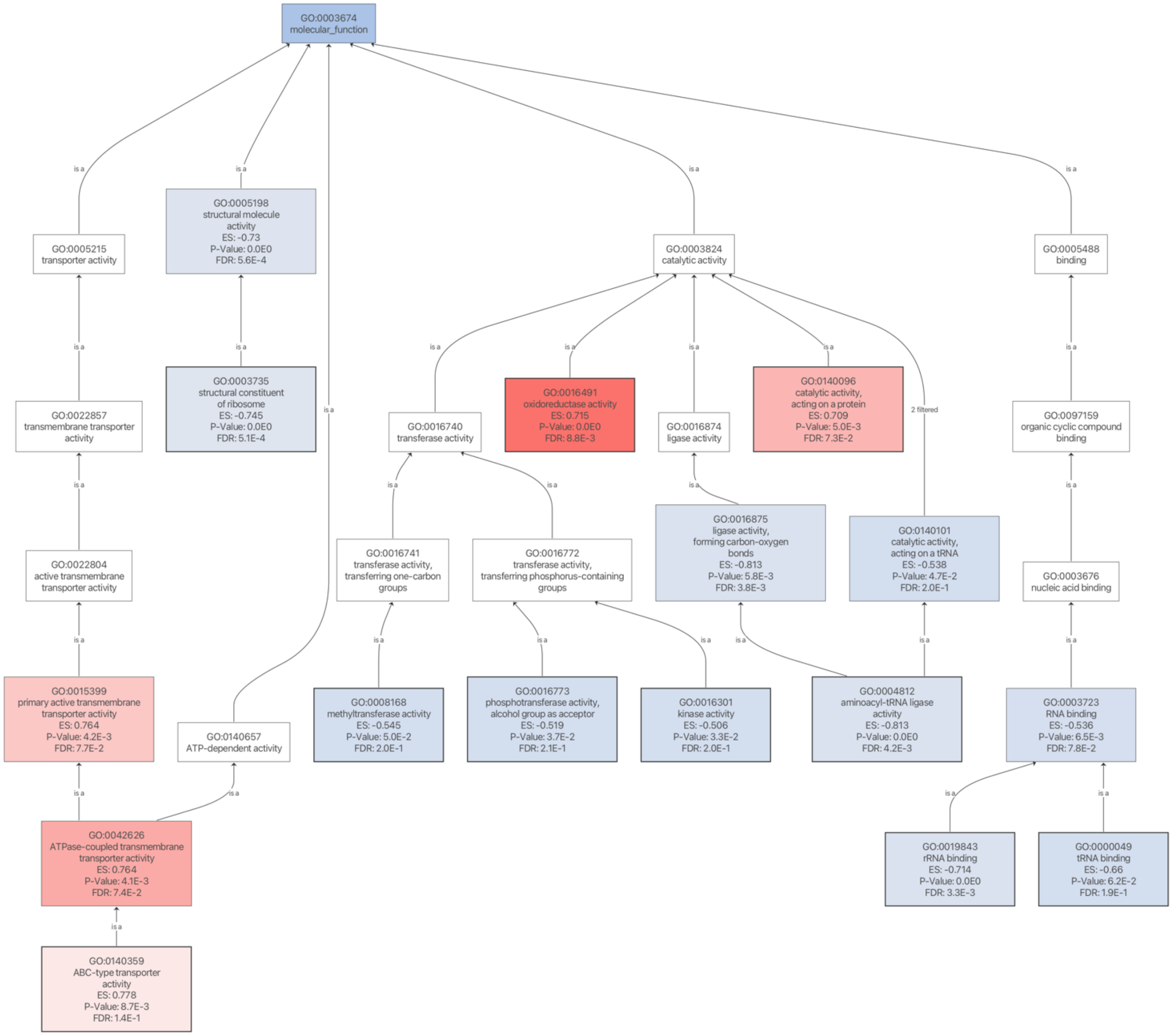
GO:0003674
molecular_function
is a
GO:0005215
transporter activity
GO:0005198
structural molecule activity
ES: -0.73
P-Value: 0.0E0
FDR: 5.6E-4
GO:0003824
catalytic activity
GO:0005488
binding
GO:0022857
transmembrane transporter activity
GO:0003735
structural constituent of ribosome
ES: -0.745
P-Value: 0.0E0
FDR: 5.1E-4
GO:0016740
transferase activity
GO:0016491
oxidoreductase activity
ES: 0.715
P-Value: 0.0E0
FDR: 8.8E-3
GO:0016874
ligase activity
GO:0140096
catalytic activity, acting on a protein
ES: 0.709
P-Value: 5.0E-3
FDR: 7.3E-2
2 filtered
GO:0097159
organic cyclic compound binding
GO:0022804
active transmembrane transporter activity
GO:0016741
transferase activity, transferring one-carbon groups
GO:0016772
transferase activity, transferring phosphorus-containing groups
GO:0016875
ligase activity, forming carbon-oxygen bonds
ES: -0.813
P-Value: 5.8E-3
FDR: 3.8E-3
GO:0140101
catalytic activity, acting on a tRNA
ES: -0.538
P-Value: 4.7E-2
FDR: 2.0E-1
GO:0003676
nucleic acid binding
GO:0015399
primary active transmembrane transporter activity
ES: 0.764
P-Value: 4.2E-3
FDR: 7.7E-2
GO:0140657
ATP-dependent activity
GO:0008168
methyltransferase activity
ES: -0.545
P-Value: 5.0E-2
FDR: 2.0E-1
GO:0016773
phosphotransferase activity, alcohol group as acceptor
ES: -0.519
P-Value: 3.7E-2
FDR: 2.1E-1
GO:0016301
kinase activity
ES: -0.506
P-Value: 3.3E-2
FDR: 2.0E-1
GO:0004812
aminoacyl-tRNA ligase activity
ES: -0.813
P-Value: 0.0E0
FDR: 4.2E-3
GO:0003723
RNA binding
ES: -0.536
P-Value: 6.5E-3
FDR: 7.8E-2
GO:0042626
ATPase-coupled transmembrane transporter activity
ES: 0.764
P-Value: 4.1E-3
FDR: 7.4E-2
GO:0019843
rRNA binding
ES: -0.714
P-Value: 0.0E0
FDR: 3.3E-3
GO:0000049
tRNA binding
ES: -0.66
P-Value: 6.2E-2
FDR: 1.9E-1
GO:0140359
ABC-type transporter activity
ES: 0.778
P-Value: 8.7E-3
FDR: 1.4E-1

**Fig. S3** Additional experiments measuring the expression of the *sodA* promoter fusion. UA159 and three mutant derivatives (*fruI, levD,* and *spxA1*), each harboring a P*sodA::gfp* fusion, were cultured in FMC containing 10 mM glucose (A), 20 mM glucose (B), 10 mM fructose (C), or 20 mM fructose (D). The relative fluorescence units (RFU) of each culture were recorded as a measure of *sodA* promoter activity, subtracted of background fluorescence from a corresponding control strain cultured under the same condition, which was of the same genetic background but Gfp-negative, and normalized against $OD_{600}$ of the cultures. Results are the average from three biological replicates, each conducted in technical duplicates.

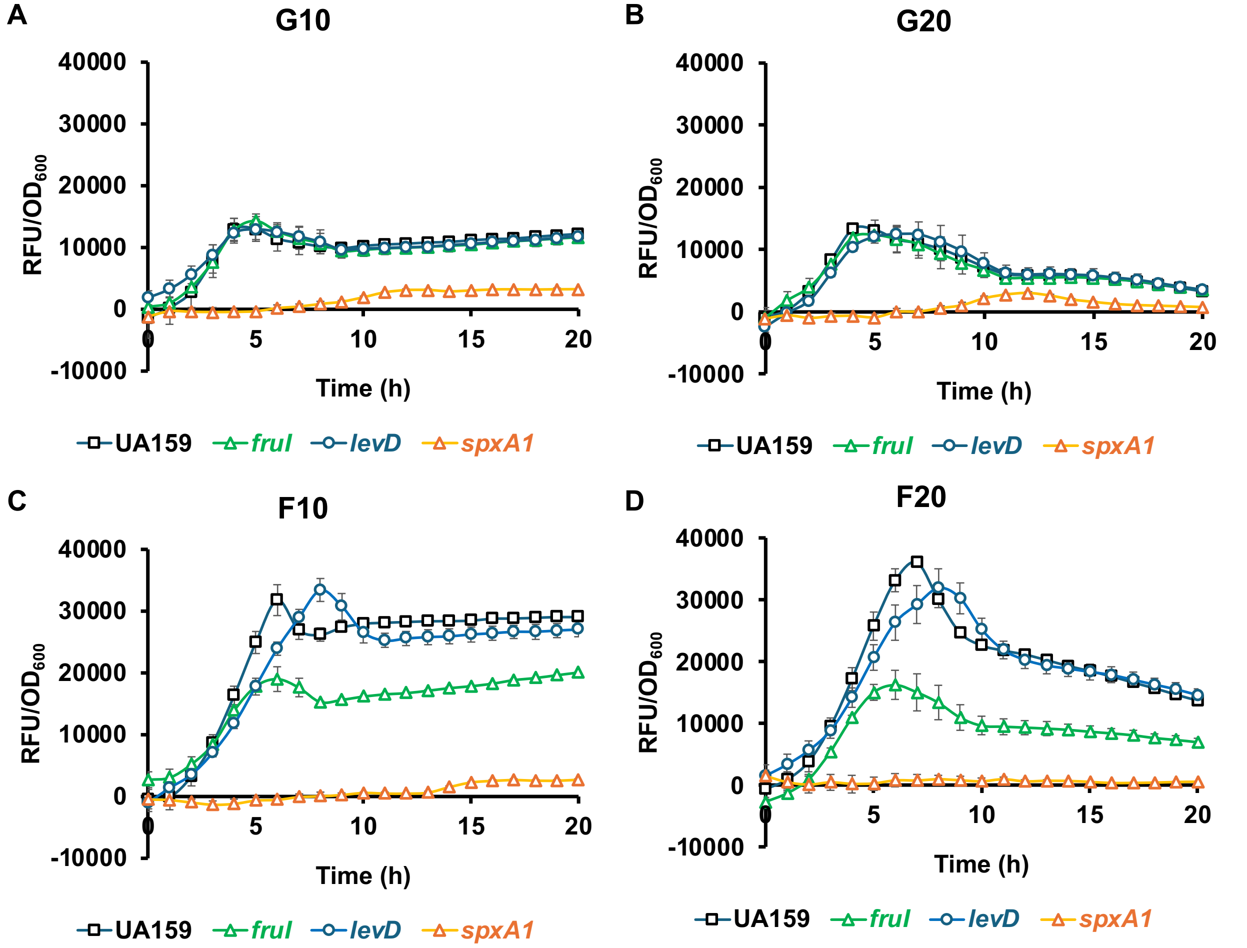

**Fig. S4** Growth of UA159/P*sodA::gfp*. BHI cultures (n = 3) of the bacterium were diluted into FMC containing 20 mM glucose (G20), 19 mM glucose plus 1 mM fructose (G19/F1), 10 mM glucose plus 10 mM fructose (G10/F10), or 20 mM glucose plus 0.5 mM $H_2O_2$. Optical density ($OD_{600}$) of these cultures was monitored for 25 h.

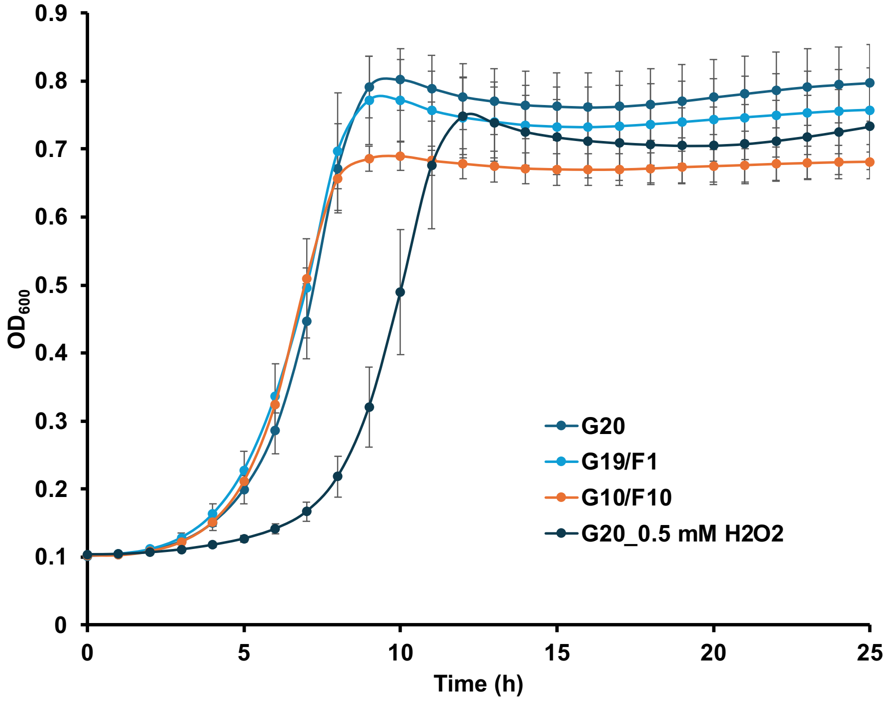

**Fig. S5** pH drop assays performed with fructose. *S. mutans* UA159 was cultivated in TV supplemented with 20 mM fructose or glucose (controls), and then subjected to glycolytic profile analysis in the presence of 50 mM fructose or glucose, respectively. pH values monitored during 1-hour period from three separate experiments (**A, B, C**) were then converted to proton concentrations, followed by statistical analysis using area-under-the-curve values (**D**). The asterisk represents statistical significance according to Student's *t* test ($P$ <0.05).

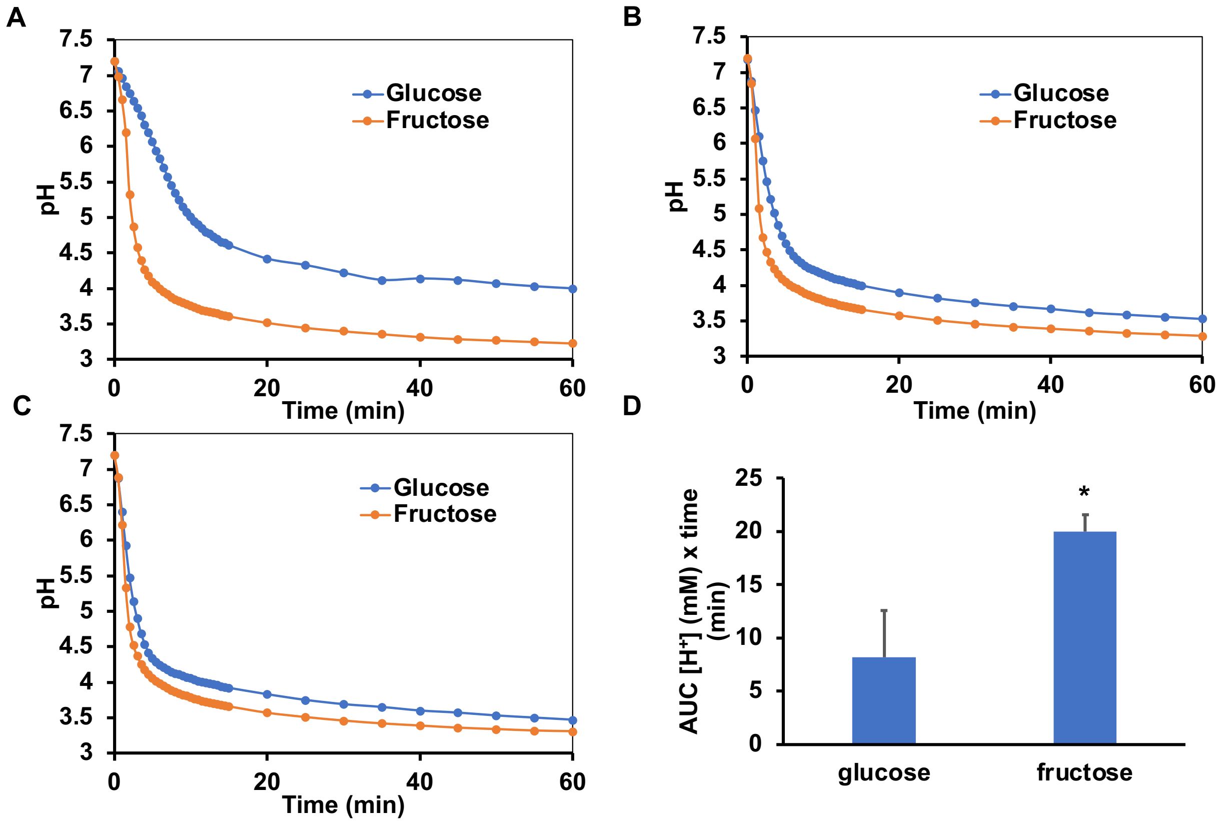

**Fig. S6** CFU recovered during competitions between *S. mutans* (SMU) and *S. sanguinis* (SSA). Exponential-phase cultures in BHI (n = 3) from both species were mixed at 1:1 (v:v) ratio, or used singly, and diluted into a FMC medium supplemented with specified amounts of carbohydrates, followed by 24 h of incubation in a 5%-$CO_2$ atmosphere. CFU enumeration was carried out to assess bacterial viability. (A) controls run with only single species in 20 mM carbohydrates. (B, C, D) mixed-species cultures enumerated at time 0 and 24 h of competition. See Fig. 7B for the final competitive indices. (E) Competition between UA159 and SK36/*spxB* in 20 mM glucose or fructose.

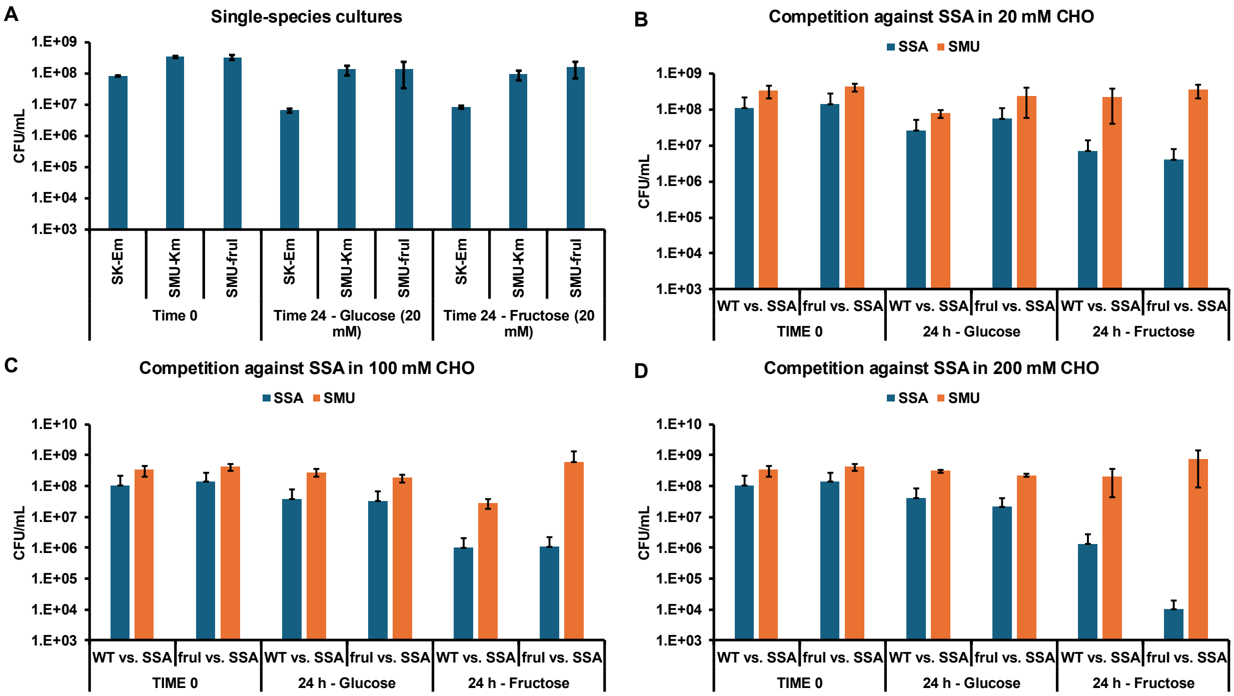

E

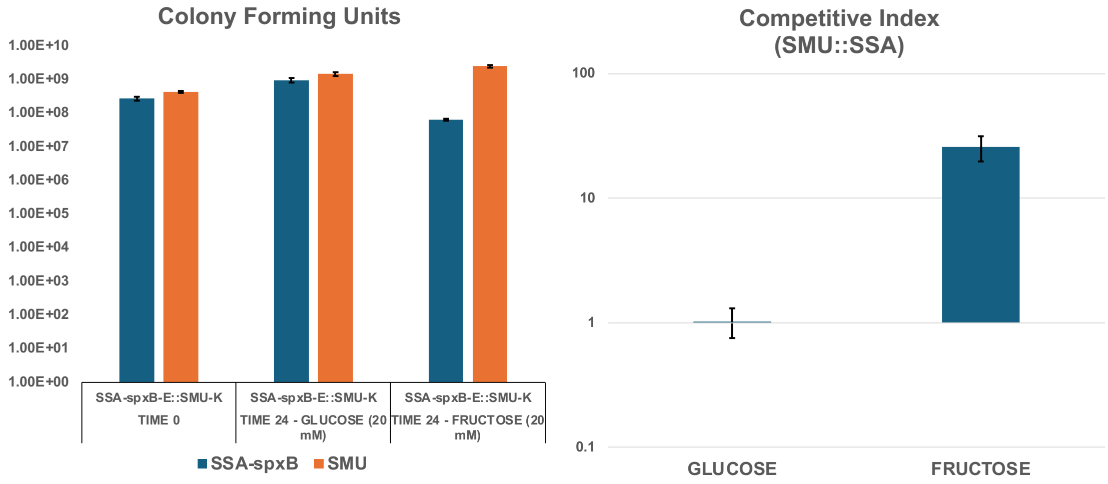
Colony Forming Units
1.00E+10
1.00E+09
1.00E+08
1.00E+07
1.00E+06
1.00E+05
1.00E+04
1.00E+03
1.00E+02
1.00E+01
1.00E+00
SSA-spxB-E::SMU-K
TIME 0
SSA-spxB-E::SMU-K
TIME 24 - GLUCOSE (20 mM)
SSA-spxB-E::SMU-K
TIME 24 - FRUCTOSE (20 mM)
SSA-spxB
SMU
Competitive Index
(SMU::SSA)
100
10
1
0.1
GLUCOSE
FRUCTOSE

1. Kajfasz JK, Ganguly T, Hardin EL, Abranches J, Lemos JA. 2017. Transcriptome responses of *Streptococcus mutans* to peroxide stress: identification of novel antioxidant pathways regulated by Spx. Sci Rep 7:16018.
